# A Dormant Microbial Component in the Development of Pre-Eclampsia

**DOI:** 10.1101/057356

**Authors:** Douglas B. Kell, Louise C. Kenny

**Affiliations:** School of Chemistry, The University of Manchester, 131, Princess St, MANCHESTER M1 7DN, Lancs, UK; The Manchester Institute of Biotechnology, The University of Manchester, 131, Princess St, MANCHESTER M1 7DN, Lancs, UK; Centre for Synthetic Biology of Fine and Speciality Chemicals, The University of Manchester, 131, Princess St, MANCHESTER M1 7DN, Lancs, UK; The Irish Centre for Fetal and Neonatal Translational Research (INFANT), University College Cork, Cork, Ireland; Department of Obstetrics and Gynecology, University College Cork, Cork, Ireland

## Abstract

Pre-eclampsia (PE) is a complex, multi-system disorder that remains a leading cause of morbidity and mortality in pregnancy. Four main classes of dysregulation accompany PE, and are widely considered to contribute to its severity. These are abnormal trophoblast invasion of the placenta, anti-angiogenic responses, oxidative stress, and inflammation. What is lacking, however, is an explanation of how these themselves are caused.

We here develop the unifying idea, and the considerable evidence for it, that the originating cause of PE (and of the four classes of dysregulation) is in fact microbial infection, that most such microbes are dormant and hence resist detection by conventional (replication-dependent) microbiology, and that by occasional resuscitation and growth it is they that are responsible for all the observable sequelae, including the continuing, chronic inflammation. In particular, bacterial products such as lipopolysaccharide (LPS), also known as endotoxin, are well known as highly inflammagenic and stimulate an innate (and possibly trained) immune response that exacerbates the inflammation further. The known need of microbes for free iron can explain the iron dysregulation that accompanies PE. We describe the main routes of infection (gut, oral, urinary tract infection) and the regularly observed presence of microbes in placental and other tissues in PE. Every known proteomic biomarker of “pre-eclampsia” that we assessed has in fact also been shown to be raised in response to infection. An infectious component to PE fulfils the Bradford Hill criteria for ascribing a disease to an environmental cause, and suggests a number of treatments, some of which have in fact been shown to be successful.

PE was classically referred to as endotoxaemia or toxaemia of pregnancy, and it is ironic that it seems that LPS and other microbial endotoxins really are involved. Overall, the recognition of an infectious component in the aetiology of PE mirrors that for ulcers and other diseases that were previously considered to lack one.

**Insight, innovation, integration:** Many descriptors of pre-eclampsia are widely accepted (e.g. abnormal trophoblast invasion, oxidative stress, inflammation and altered immune response, and anti-angiogenic responses). However, without knowing what causes them, they do not explain the syndrome. The Biological Insight of this manuscript is that there is considerable evidence to the effect that each of these phenomena (hence PE) are caused by the resuscitation of dormant bacteria that shed (known and potent) inflammagens such as LPS, often as a consequence of iron availability. PE is thus seen as a milder form of sepsis. The Technological Innovations come from the use of molecular markers (of microbes and omics more generally, as well as novel markers of coagulopathies) to measure this. The Benefit of Integration comes from bringing together a huge number of disparate observations into a unifying theme.

## Introduction

**Pre-eclampsia.** Pre-eclampsia (PE) is a multi-system disorder of pregnancy, characterised and indeed defined by the presence of hypertension after 20 weeks’ gestation and before the onset of labour, or postpartum, with either proteinuria or any multisystem complication [1–8]. It is a common condition, affecting some 3-5% of nulliparous pregnant women [7; 9]and is characterised by high mortality levels [10–13]. There is no known cure other than delivery, and consequently PE also causes significant perinatal morbidity and mortality secondary to iatrogenic prematurity. There are a variety of known risk factors (Table 1), that may be of use in predicting a greater likelihood of developing PE, albeit there are so many, with only very modest correlations, that early-stage (especially first-trimester) prediction of late-stage PE remains very difficult [7; 14; 15].

It is striking that most of the ‘risk factors’ of Table 1 are in fact risk factors for <u>multiple</u> vascular or metabolic diseases, i.e. they merely <u>pre-dispose</u> the individual to a greater likelihood of manifesting the disease or syndrome (in this case PE). Indeed, some of them are diseases. This would be consistent with the well-known comorbidities e.g. between PE and later cardiovascular disease (e.g. [16–26]), between PE and intracerebral haemorrhage during pregnancy (OR 10.39 [27]), and between PE and stroke post-partum [28; 29]. The penultimate row of Table 1 lists a series of diseases that amount to comorbidities, although our interest was piqued by the observation that one third of patients with anti-phospholipid syndrome have PE, and infectious agents with known cross-reacting antigens are certainly one original (external) source of the triggers that cause the anti-phospholipid antibodies [30–33] (and see below). Similarly, in the case of urinary tract infection, the ‘risk’ factor is a genuine external trigger, a point (following the call [34] by Mignini and colleagues for systematic reviews) that we shall expand on considerably here.

**Table 1.**
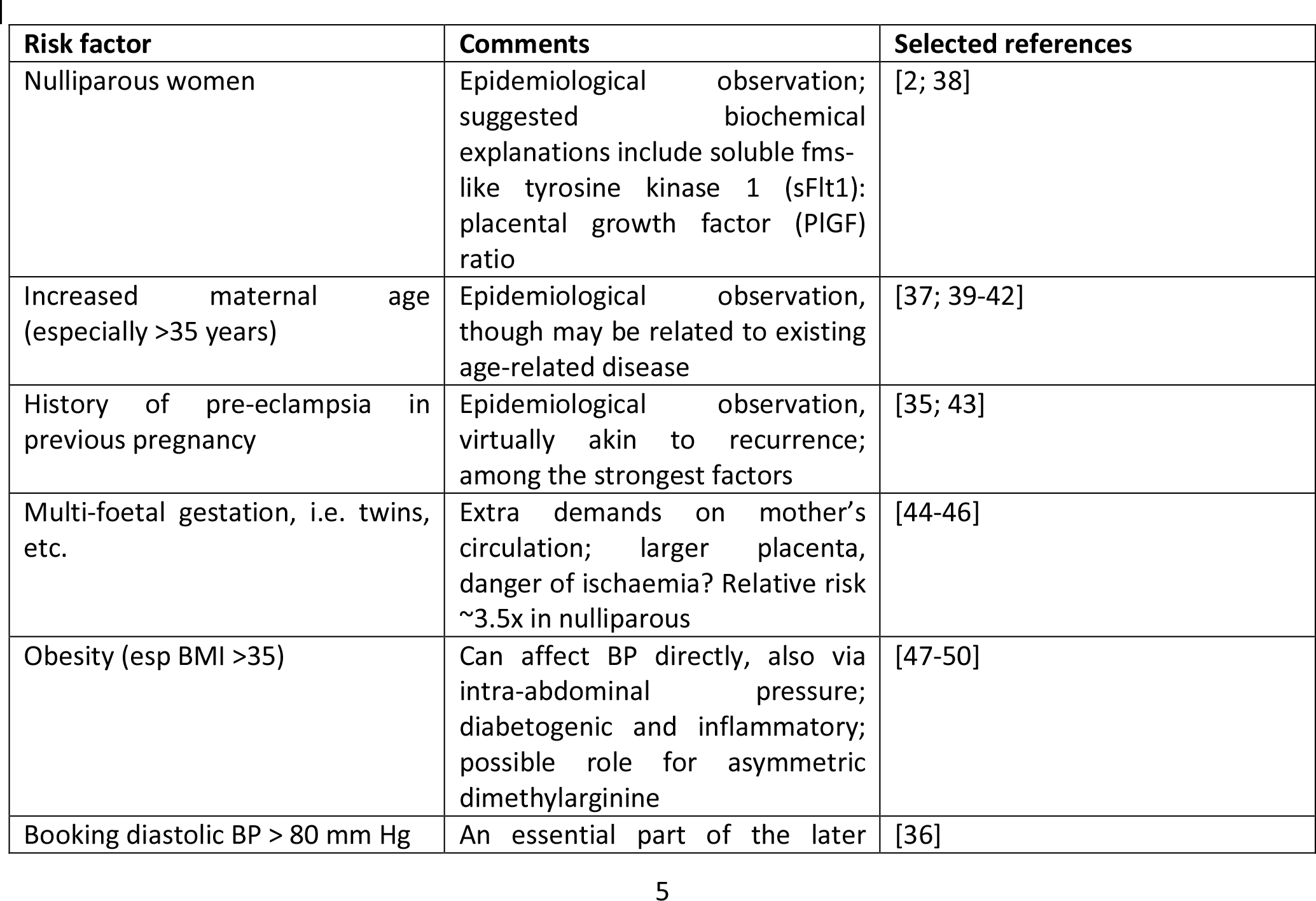
Some known risk factors for pre-eclampsia (based in part on [2; 6; 35-37]). See also http://bestpractice.bmj.com/best-practice/monograph/326/diagnosis.html. Note that most of these are risk factors that might and do pre-dispose for other diseases (or are themselves diseases).

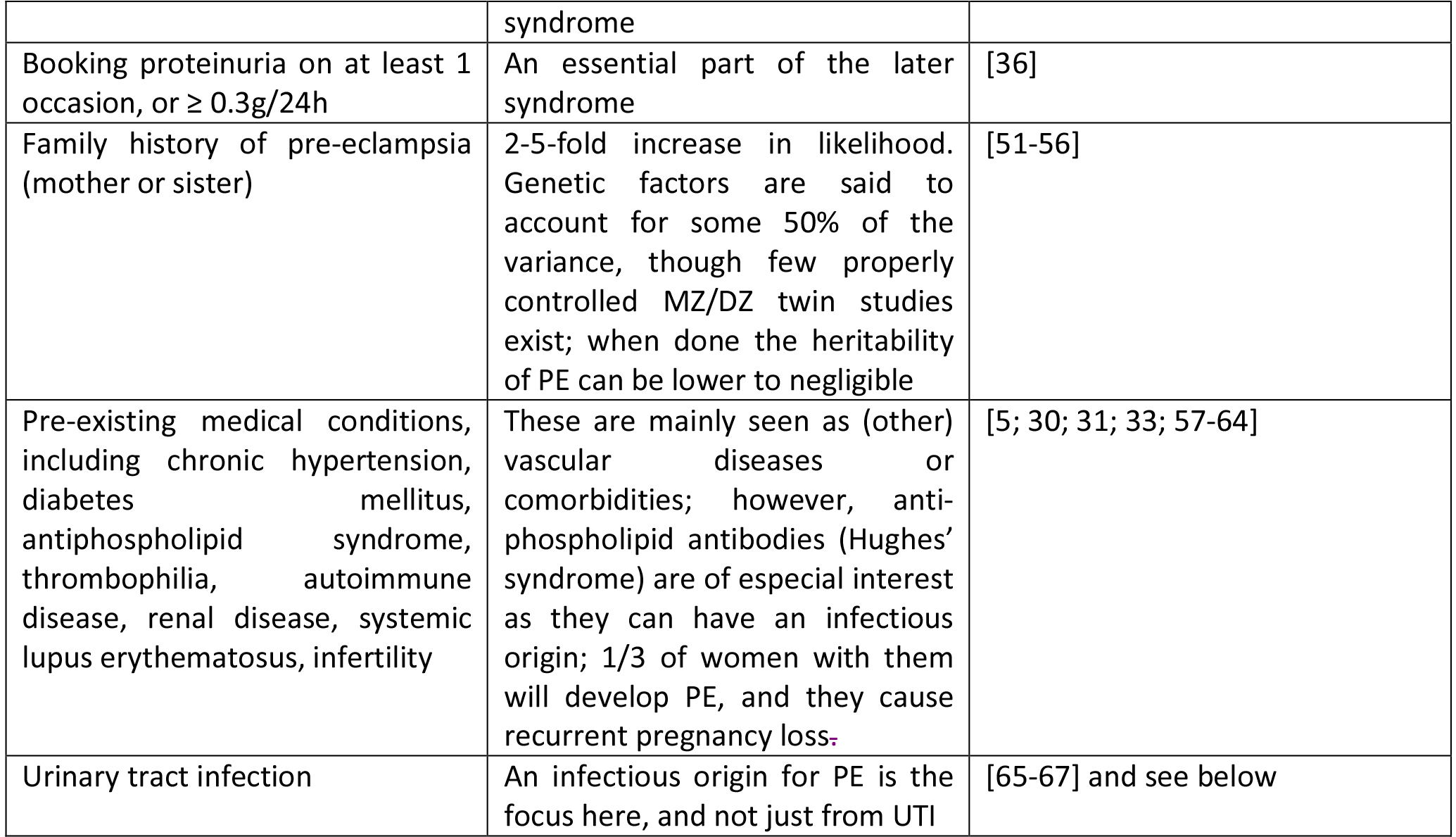

In recent decades, intense investigation has led to the development of a two-stage aetiological model for pre-eclampsia, first proposed by Redman and colleagues [68], in which inadequate remodelling of the spiral arteries in early gestation results in poor placental development (stage one) and the resultant ischaemia/re-perfusion injury and oxidative stress eventually leads to maternal vascular endothelial cell dysfunction and the maternal manifestations of the disease (stage 2) [68–72]. However, many clinical inconsistencies challenge the simplicity of this model. For example, whilst the association between poor placentation and pre-eclampsia is well established, it is not specific. Poor placentation and foetal growth restriction (FGR) frequently present without maternal signs of pre-eclampsia. Moreover, FGR is not a consistent feature of pre-eclampsia. Whilst it is commonly seen in pre-eclampsia presenting at earlier gestations, in pre-eclampsia presenting at term, neonates are not growth restricted and may even be large for dates [73].

Thus, the two-stage model has been further refined by Roberts and others [72; 74; 75] to take into account the heterogeneous nature of pre-eclampsia and the varying contribution from mother and infant to the disorder. We now appreciate that normal pregnancy is characterised by a low-grade systemic inflammatory response and specific metabolic changes, and that virtually all of the features of normal pregnancy are simply exaggerated in pre-eclampsia [76–78]. There is also widespread acceptance that maternal constitutional and environmental factors (such as obesity) can interact to modulate the risk of preeclampsia. Thus, with profoundly reduced placental perfusion (or significant ‘placental loading’), the generation of Stage 2 may require very little contribution from the mother to provide sufficient stress to elicit the maternal syndrome. In this setting, almost any woman will develop pre-eclampsia. Conversely, the woman with extensive predisposing constitutional sensitivity could develop pre-eclampsia with very little reduced perfusion, or minimal ‘placental loading’. As with many complex disorders, multiple factors can affect disease development positively or negatively, with a convenient representation of the two main negative sources (foetal and maternal) being that of a see-saw [79], as in Fig 1.

**Figure 1.**
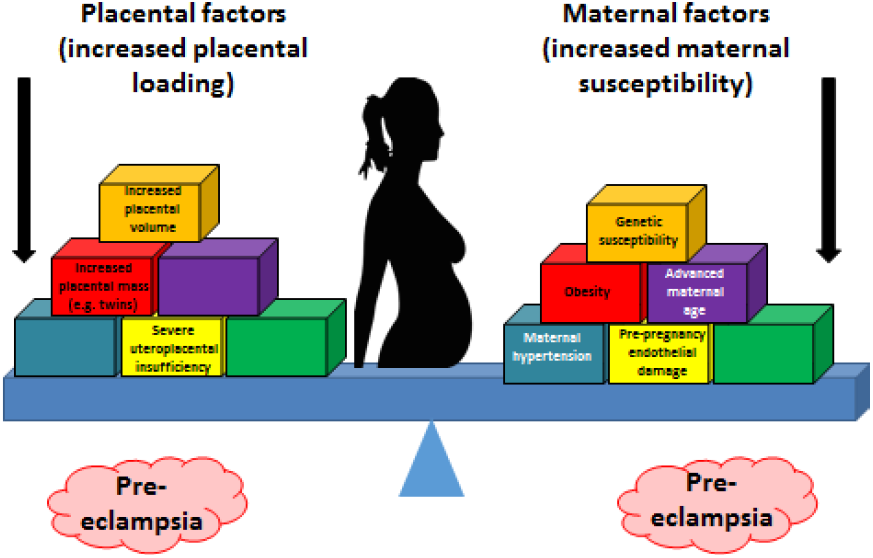
Two main sources (foetal and maternal) can drive a pregnancy towards pre-eclampsia.

Whilst this explains the inconsistencies of the two-stage model, the precise mechanisms 1) underlying the initial poor placentation and 2) linking placental stress and the maternal syndrome have still not been fully elucidated.

Much recent research in pre-eclampsia has focused on various angiogenic factors, including the pro-angiogenic factors vascular endothelial growth factor (VEGF) and placental growth factor (PlGF) and the two anti-angiogenic proteins, soluble endoglin (sEng) and soluble fms-like tyrosine kinase 1 (sFlt). Recent data suggest that alterations in circulating angiogenic factors play a pathogenic role in pre-eclampsia. These angiogenic factors tightly regulate angiogenesis and are also essential for maintenance of normal vessel health. Consequently, the synthesis and action of these factors and their receptors in the uterine bed and placenta are essential for normal placental development and pregnancy [80; 81]. In pre-eclampsia, increased levels of the anti-angiogenic sFlt-1 and sEng trap circulating VEGF, PlGF and transforming growth factor P (TGFP) respectively. A myriad of data supports the idea that circulating levels of these factors alone, or in combination, can be used to predict pre-eclampsia [82; 83] (and see below under PE biomarkers), but in line with the heterogeneous nature of pre-eclampsia, the data are somewhat inconsistent and their performance as biomarkers seems limited to disease with significant placental loading [7]. Therefore, angiogenic dysregulation would appear unlikely to be the sole link between the stressed placenta and endothelial dysfunction and the clinical manifestations of the disease.

Notwithstanding these many inconsistencies, the central role of the placenta as a source of ‘toxin’, in a condition regarded, and indeed often named, as ‘toxaemia of pregnancy’ [84–86] cannot be refuted. The uncertainty regarding the nature of the toxin(s) continues, and other placental sources of endothelial dysfunction include syncytiotrophoblast basement membrane fragments (STBM) [87] and endothelial progenitor cells (EPC) [88]; an increase of reactive oxygen species over scavenging by anti-oxidants [89; 90] has also been promoted.

The Bradford Hill criteria for causation of a disease Y by an environmental factor X [91] are as follows:

(1) strength of association between X and Y; (2) consistency of association between X and Y; (3) specificity of association between X and Y; (4) experiments verify the relationship between X and Y; (5) modification of X alters the occurrence of Y; (6) biologically plausible cause and effect relationship.

In general terms [92], if we see that two things (A and B) co-vary in different circumstances, we might infer that A causes B, that B causes A, or that something else (C) causes both B and A, whether in series or parallel. To disentangle temporal relations requires a longitudinal study. The job of the systems biologist doing systems medicine is to uncover the chief actors and the means by which they interact [93], in this way fulfilling the Bradford Hill postulates, a topic to which we shall return at the end.

In infection microbiology, and long predating the Bradford Hill criteria, the essentially equivalent metrics are known (widely, but somewhat inaccurately [94]) as the Koch or Henle-Koch postulates (i.e. criteria). They involve assessing the correlation of a culturable organism with the presence of a disease, the cure of the disease (and its symptoms) upon removal of the organism, and the development of the disease with (re)inoculation of the organism. They are of great historical importance, but present us with three main difficulties here. The first is that we cannot apply the third of them to humans for obvious ethical reasons. The second (see also below) and related one is that we cannot usefully apply them in animal models because none of the existing models recapitulates human pre-eclampsia well. Finally, as widely recognised [94–101], they cannot be straightforwardly applied when dealing with dormant bacteria or bacteria that are otherwise refractory to culture.

Our solution to this is twofold: (i) we can assess the first two using molecular methods if culturing does not work, and (ii) we exploit the philosophy of science principle known as ‘coherence’ [102–106]. This states that if a series of ostensibly unrelated findings are brought together into a self-consistent narrative, that narrative is thereby strengthened. Our systems approach purposely represents a ‘coherence’ in the sense given.

**Figure 2.**
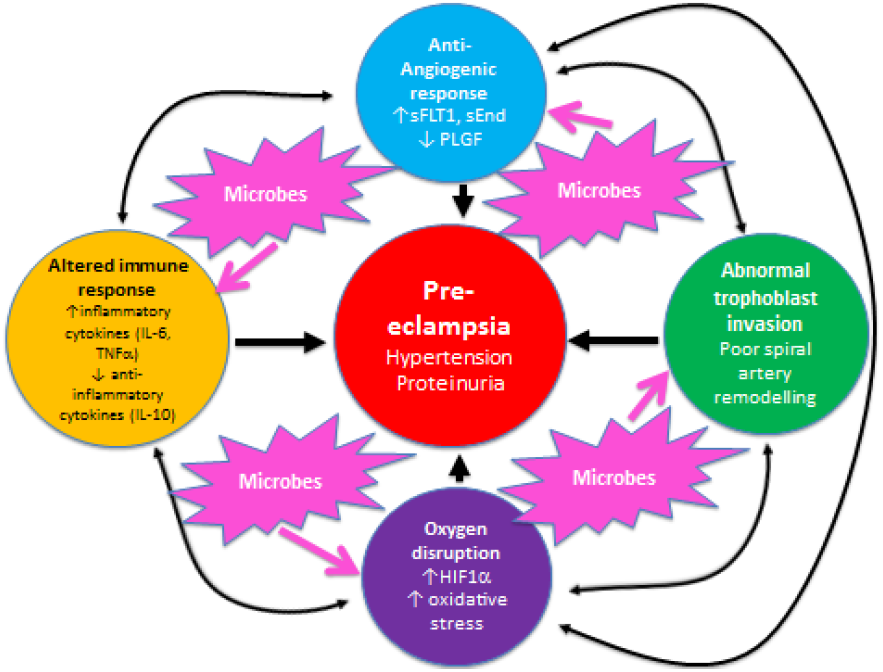
There are four main ‘causes’ of pre-eclampsia, represented by the coloured outer circles, and these too can interact with each other. That part of the figure is redrawn from [107]. In addition, we note here, as the theme of this review, that microbes can themselves cause each of the features in the outer coloured circles to manifest.

Overall, known biochemical associations with PE come into four main categories, viz. abnormal trophoblast invasion, oxidative stress, inflammation and altered immune response, and anti-angiogenic responses (Fig 2). Each of these can contribute directly to PE, and although they can interact with each other (black arrows), no external or causal source is apparent. Fig 2 has been redrawn from a very nice review by Pennington and colleagues [107], which indicates four main generally accepted ‘causes’ (or at least accompaniments) of PE as the four outer coloured circles. As illustrated with the black two-way arrows, many of these also interact with each other. What is missing, in a sense, is then what causes <u>these</u> causes, and that is the nub of our argument here. Since we now know (and describe below) that microbes can affect <u>each</u> of these four general mechanisms, we have added these routes to Fig 1 (using pink arrows) where dormant, resuscitating or growing microbes are known to contribute.

In a similar vein, Magee and colleagues [108] have nicely set down their related analysis of the causes and consequences of PE, with a central focus (redrawn in Fig 3) on endothelial cell activation. While bearing much similarity in terms of overall content to the analysis of Pennington and colleagues [107], and ours above, it again lacks a microbial or infection component as a <u>causative</u> element, but importantly does note that infection and/or inflammation can serve to lower the threshold for PE in cases of inadequate placentation. In our view microbes can also enter following normal placentation if their dormant microbiome begins to wake up and/or to shed inflammagens.

**Figure 3.**
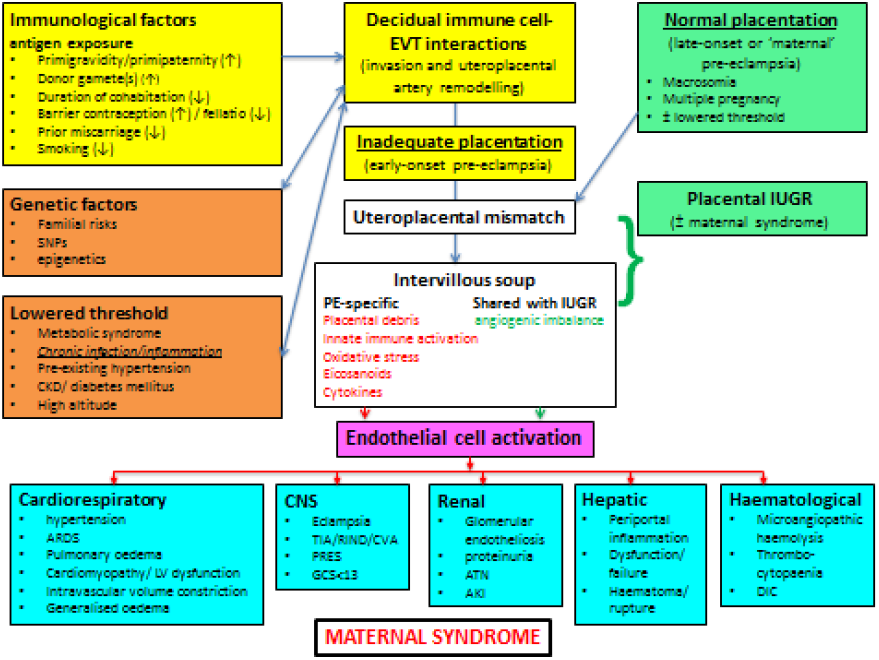
Another detailed representation of factors known to cause or accompany PE, redrawn from [108]

**Heritability**. The question of the extent of heritability of PE (susceptibility) is of interest. Although this seems to vary widely in different studies (Table 1), a number of candidate gene studies [54; 109–112] imply that a susceptibility to PE is at least partly heritable, consistent with the variance in all the other ‘risk factors’ of Table 1 (and see [5]). As with all the other gene-association studies where phenotypic (‘lifestyle’) information is absent [113–115], it is not possible to ascribe the heritability to genetics alone, as opposed to an interaction of a genetic susceptibility (e.g. in the HLA system) with environmental factors [111], such as cytomegalovirus infection [116].

**Inflammation**. Pre-eclampsia is accompanied by oxidative stress [117] and inflammation, and thus shares a set of observable properties with many other (and hence related) inflammatory diseases, be they vascular (e.g. atherosclerosis), neurodegenerative (e.g. Alzheimer’s, Parkinson’s), or ‘metabolic’ (type 1 and 2 diabetes). It is thus at least plausible that they share some common aetiologies, as we argue here, and that knowledge of the aetiology of those diseases may give us useful clues for PE.

As well as raised levels of inflammatory cytokines, that constitute virtually a circular definition of inflammation, we and others have noted that <u>all</u> of these diseases are accompanied by dysregulation of iron metabolism [79; 118; 119]], hypercoagulability and hypofibrinolysis [120; 121], blood microparticles [119], and changes in the morphology of fibrin fibres (e.g. [122–127]) and of erythrocytes (e.g. [120; 125–130]).

In addition, we and others have recognised the extensive evidence for the role of a dormant blood and/or tissue microbiome in these [131–136] and related [137–140] diseases, coupled in part to the shedding of highly inflammagenic bacterial components such as Gram-negative lipopolysaccharides (LPS) and their Gram-positive cell wall equivalents such as lipoteichoic acids [141]. (We shall often use the term ‘LPS’ as a ‘shorthand’, to be illustrative of all of these kinds of highly inflammagenic molecules.)

The purpose of the present review, outlined as a ‘mind map’ in Fig 4, is thus to summarise the detailed and specific lines of evidence suggesting a very important role of a dormant microbial component in the aetiology of pre-eclampsia (and see also [131]). To do this, we must start by rehearsing what is meant by microbial dormancy.

**Figure 4.**
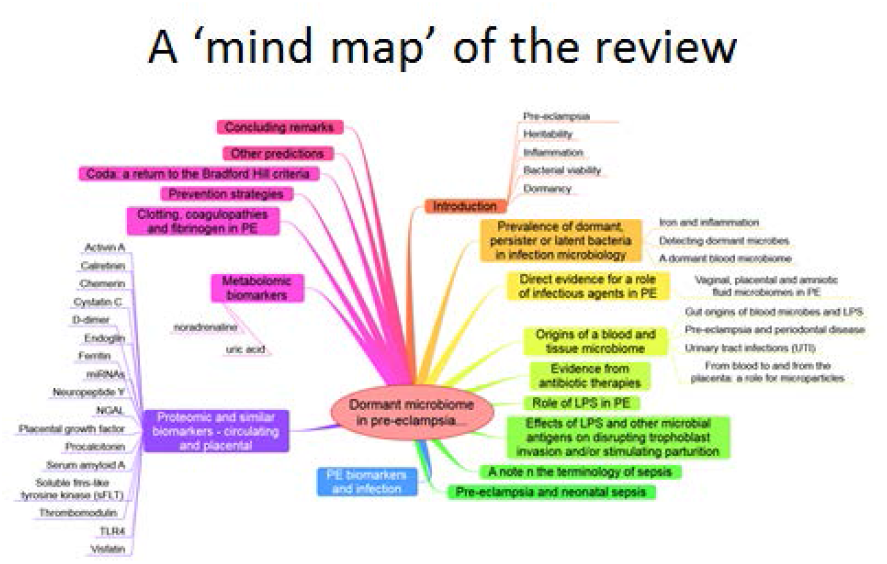
A mind map of the overall structure of the review

**Bacterial viability**. In microbiology, we usually consider microbes as being in one of three ‘physiological macrostates’ (Fig 5). The definition of a ‘viable’ bacterium is normally based on its ability to replicate, i.e. ‘viability’ = culturability [142–144]. In this sense, classical microbiology has barely changed since the time of Robert Koch, with the presence of a ‘viable’ microorganism in a sample being assessed via its ability to form a visible colony on an agar plate containing suitable nutrients. However, it is well known, especially in environmental microbiology (‘the great plate count anomaly’ [145]), that only a small percentage of cells observable microscopically is typically culturable on agar plates. In principle this could be because they are or were ‘irreversibly’ non-culturable (operationally ‘dead’), or because our culture media either kill them [146] or such media lack nutrients or signalling molecules necessary for their regrowth [147; 148] from an otherwise dormant state [149; 150]. Those statements are true even for microbes that appear in culture collections and (whose growth requirements) would be regarded as ‘known’.

**Figure 5.**
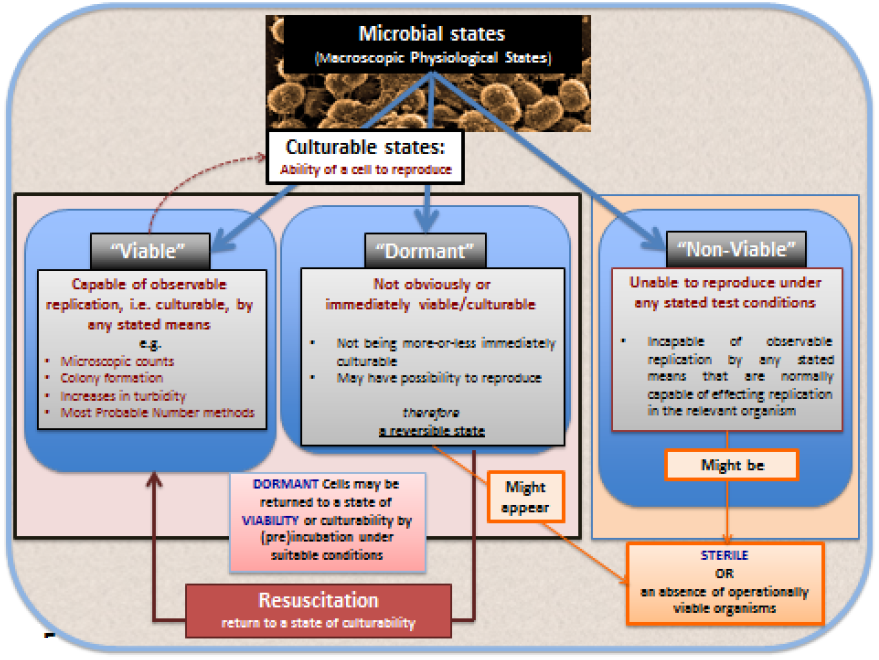
The chief physiological macrostates exhibited by microorganisms

However, it is common enough in clinical microbiology that we detect the existence or presence of ‘novel’ microbial pathogens with obscure growth requirements before we learn to culture them; this is precisely what happened in the case of *Legionella pneumophila* [151–154], *Tropheryma whipplei* (Whipple’s disease [155; 156]), and *Coxiella burnetii* (the causative agent of Q fever [157; 158]). Even *Helicobacter pylori* was finally brought into culture on agar plates only because an unusually long Easter holiday break meant that the plates were incubated for an extended period of five days (rather than the normal two) before being thrown out [159; 160]! Consequently, there is ample precedent for the presence of ‘invisible’ microbes to go unremarked before they are discovered as the true cause of a supposedly non-infectious disease, even when they are perfectly viable (culturable) according to standard analyses.

**Dormancy** for a microbe is defined operationally as a state, commonly of low metabolic activity, in which the organism appears not to be viable in that it is unable to form a colony but where it is not dead in that it may revert to a state in which it can do so, via a process known as resuscitation [149; 150]. However, an important issue (and see above) is that <u>dormant</u> bacteria do not typically fulfil the Koch-Henle postulates [94; 96–98], and in order for them to do so it is necessary that they be grown or resuscitated. This is precisely what was famously done by Barry Marshall and Robin Warren when they showed that the <u>**supposedly** non-infectious</u> disease of gastric ulcers was in fact caused by a ‘novel’ organism called <u>Helicobacter pylori</u> [161; 162]. One of the present authors showed in laboratory cultures of actinobacteria that these too could enter a state of true dormancy [163; 164] (as is well known for <u>Mycobacterium tuberculosis</u>, e.g. [165–169]), and could be resuscitated by a secreted growth factor called Rpf [170–174]. This RPF family has a very highly conserved motif that is extremely immunogenic [175; 176], and it is presently under trials as a vaccine against *M. bovis.*

**Prevalence of dormant, persistent or latent bacteria in infection microbiology**. It is worth stressing here that the presence of dormant or latent bacteria in <u>infection microbiology</u> is well established; one third of humans carry dormant *Mycobacterium tuberculosis* (e.g. [165; 177–180]), most without reactivation, while probably 50-100% are infected with *H. pylori,* most without getting ulcers or worse [181; 182]. As with the risk factors in Table 1, the organisms are merely or equivalently ‘risk factors’ for those infectious diseases and are effectively seen as causative only when the disease is actually manifest.

In a similar vein, so-called persisters are <u>phenotypic</u> variants of infectious microbes that resist antibiotics and can effectively lie in hiding to resuscitate subsequently. This too is very well established (e.g. [132; 183–196]). In many cases they can hide intracellularly [197], where antibiotics often penetrate poorly [198] because the necessary transporters [199–202] are absent. This effectively provides for reservoirs of reinfection, e.g. for *Staphylococcus aureus* [203], *Bartonella* spp [204] and-most pertinently here-for the *Escherichia coli* involved in urinary tract (re)infection [205–208]. The same intracellular persistence is true for parasites such as *Toxoplasma gondii* [209].

Thus, the main point of the extensive prevalence of microbial dormancy and persistence is that microbes can appear to be absent when they are in fact present at high concentrations. This is true not only in cases where infection is recognised as the cause of disease but, as we here argue, such microbes may be an important part of diseases presently thought to lack an infectious component.

**Iron and inflammation**. It is well known that (with the possible exception of *Borrelia* [210; 211]) a lack of free iron normally limits microbial growth *in vivo* (e.g. [212–236]), and we have reviewe—d previously [79; 118; 119]] the very clear iron dysregulation accompanying pre-eclampsia (e.g. [84; 237–249]).

This has led to the recognition [121; 132; 134] that the source of the <u>continuing</u> inflammation might be iron-based resuscitation of dormant microbes that could release well-known and highly potent inflammagens such as lipopolysaccharide (LPS). Indeed, we have shown that absolutely tiny (highly substoichiometric) amounts of LPS can have a massive effect on the blood clotting process [250], potentially inducing β-amyloid formation directly [251; 252] (something, interestingly, that can be mimicked in liquid crystals [253; 254]). The overall series of interactions envisaged (see also [132]) is shown in Fig 6.

**Figure 6.**
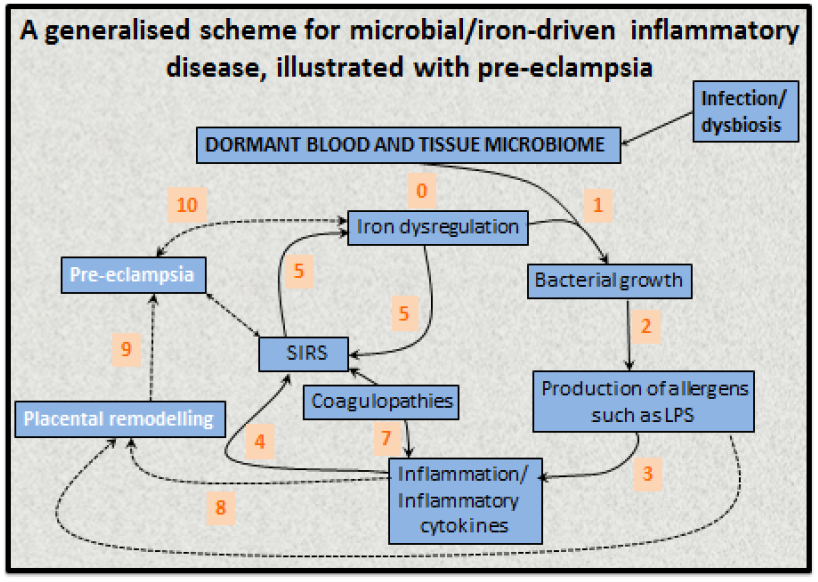
An 11-stage systems biology model of the factors that we consider cause initially formant microbes to manifest the symptoms (and disease) of pre-eclampsia

**Detecting dormant microbes**. By definition, dormant bacteria escape detection by classical methods of assessing viability that involve replication on agar plates. Other growth-associated methods include measurements involving changes in turbidity [255], including an important but now rather uncommon technique referred to as the ‘most probable number’ (MPN). The MPN involves diluting samples serially and assessing by turbidity changes the presence of growth/no growth. Look-up tables based on Poisson statistics enable estimation of the number of cells or propagules that were present. A particular virtue is that they allow dormant and ‘initially viable’ cells to be discriminated via ‘dilution to extinction’ [164], thereby avoiding many artefacts [150]. As mentioned above, preincubation in a weak nutrient broth [164; 256] was instrumental in allowing the discovery [170] of an autocrine ‘wake-up’ molecule necessary for the growth of many actinobacteria.

Other more classical means of detecting microbes, but not whether they were <u>culturable</u>, involved microscopy [183; 257-260] or flow cytometry [261] with or without various stains that reflected the presence or otherwise of an intact cell wall/membrane [163; 262-269]. These stains are sometimes referred to as ‘viability’ stains, but this is erroneous as they do not measure ‘culturability’. Readers may also come upon the term ‘viable-but-not-culturable’ however, since viable = culturable, this is an oxymoron that we suggest is best avoided [150]. Other methods involved measurement of microbial products, e.g. CO_2_ [270; 271], or changes in the conductivity or impedance of the growth medium [255; 272–274].

Most importantly, however, dormant (as well as culturable) cells may be detected by molecular means, nowadays most commonly through PCR and/or sequencing of the DNA encoding their small subunit ribosomal RNA (colloquially ‘16S’) [275–289] or other suitable genes. It is clear that such methods will have a major role to play in detecting, identifying and quantifying the kinds of microbes that we argue lie at the heart of PE aetiology.

**A dormant blood microbiome**. Of course actual bacteraemia, the presence of replicable bacteria in blood, is highly life-threatening [290], but-as emphasised-viability assays do not detect <u>dormant</u> bacteria. When molecular detection methods are applied to human blood, it turns out that blood does indeed harbour <u>a great many</u> dormant bacteria (e.g. [291–301]); they may also be detected ultramicroscopically (e.g. [132–134; 183; 259; 292; 302]) or by flow cytometry [303], and dormant blood and tissue microbes probably underpin a great many chronic, inflammatory diseases normally considered to lack a microbial component [132–134; 137–140; 183; 259; 260; 294; 304–313]. Multiple arguments serve to exclude ‘contaminants’ as the source of the bacterial DNA [134]: 1. There are significant differences between the blood microbiomes of individuals harbouring disease states and nominally healthy controls, despite the fact that samples are treated identically; 2. The morphological type of organism (e.g. coccus vs bacillus) seems to be characteristic of particular diseases; 3. In many cases relevant organisms lurk intracellularly, which is hard to explain by contamination; 4. There are just too many diseases where bacteria have been found to play a role in the pathogenesis, that all of them may be caused by contamination; 5. The actual numbers of cells involved seem far too great to be explicable by contamination; given that blood contains ~5.10^9^ erythrocytes.mL^−1^, if there was just one bacterial cell per 50,000 erythrocytes this will equate to 10^5^ bacteria.mL^−1^. These are big numbers, and if the cells were culturable, that number of cells would be the same as that ordinarily defining bacteriuria.

A recent study by Damgaard and colleagues [298] is of particular interest here. Recognising the strong mismatch between the likelihood of an infection post-transfusion (very high [298]) and the likelihood of detecting culturable microbes in blood bank units (negligible, ca 0.1%) [298; 314], Damgaard *et al* reasoned that our methods of detecting and culturing these microbes might be the problem. Certainly, taking cells from a cooled blood bag and placing them onto an agar plate at room temperature that is directly exposed to atmospheric levels of gaseous O_2_ is a huge stress leading to the production of ‘reactive oxygen species’ [118; 315], that might plausibly kill any dormant, injured, or even viable microbes. Thus they incubated samples from blood on a rich medium (trypticase soy agar) for a full week, both aerobically and anaerobically. Subsequent PCR and sequencing allowed them to identify specific microbes in some 35-53% of the samples. Thus, very careful methods need to be deployed to help resuscitate bacteria from physiological states that normally resist culture, even when those bacteria are well-established species. This is very much beginning to happen in environmental microbiology (e.g. [147; 316–318]), and such organisms are rightly seen as important sources of novel bioactives [319; 320].

As reviewed previously [132–136], the chief sources of these blood microbes are the gut microbiome, the oral microbiome (periodontitis [321]), and via urinary tract infections. Consequently, if we are to argue that there is indeed a microbial component to pre-eclampsia, we should expect to see some literature evidence for it [66; 67; 131; 322–324]. In what follows we shall rehearse the fact that it is voluminous.

## Direct evidence for a role of infectious agents in PE

Although we recognise that many of the more molecular methods cannot distinguish culturable from dormant microbes, quite a number of studies have explicitly identified infection as a cause of PE (Table 2). The commonest microbe seems to be *H. pylori*; while it is most famously associated with gastric ulcers [161; 162; 325], there are many other extragastric manifestations (e.g. [326–334]). The Odds Ratio of no less than 26 in PE vs controls when the strains can produce CagA antigens is especially striking, not least because it provides a mechanistic link to poor trophoblast invasion via a mechanism involving host antibodies to CagA cross-reacting with trophoblasts [335; 336], and circulating [337] in microparticles [338] or endosomes [339; 340].

**Table 2.**
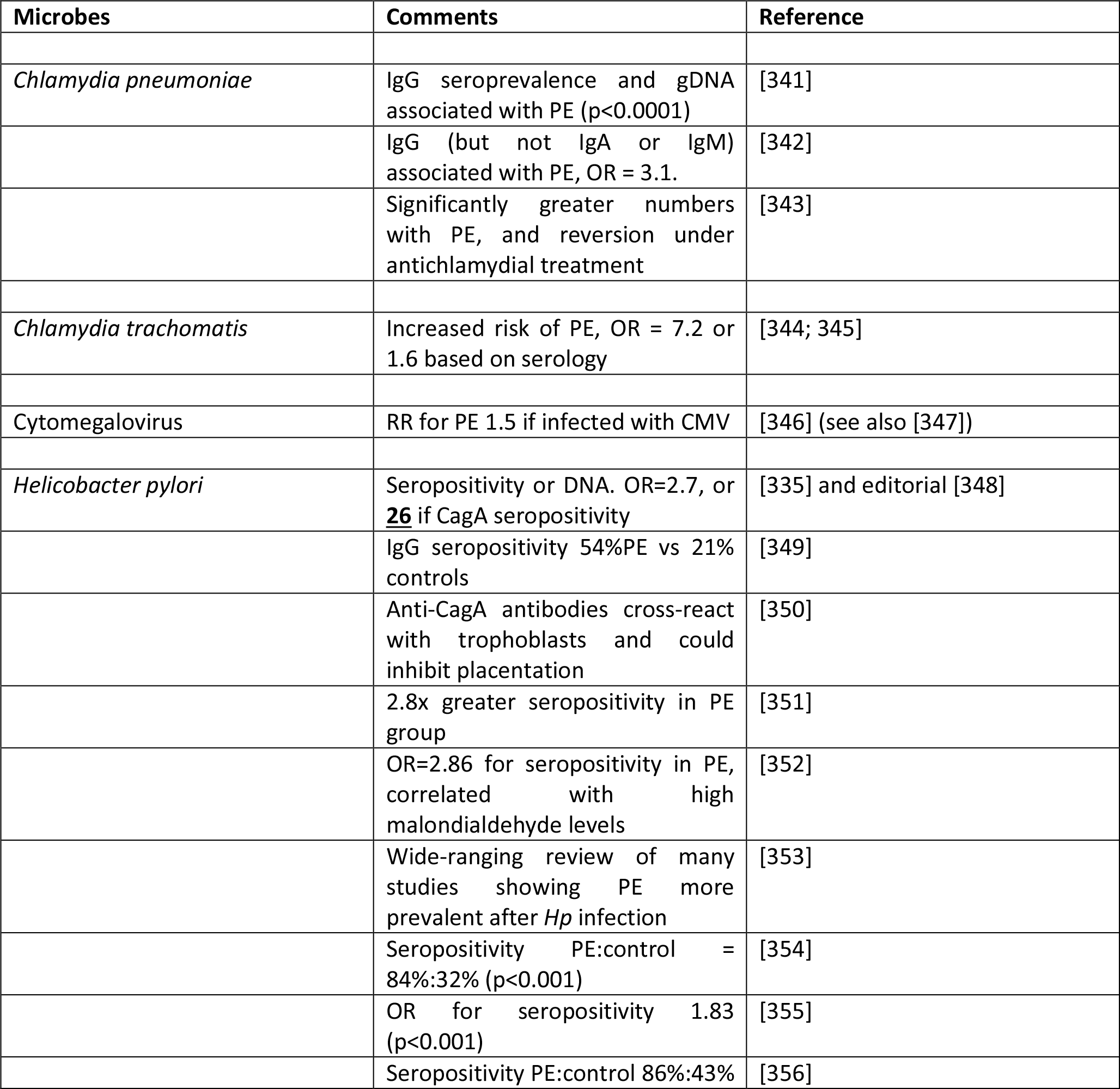
Many studies have identified a much greater prevalence of infectious agents in the blood or urine of those exhibiting PE than in matched controls

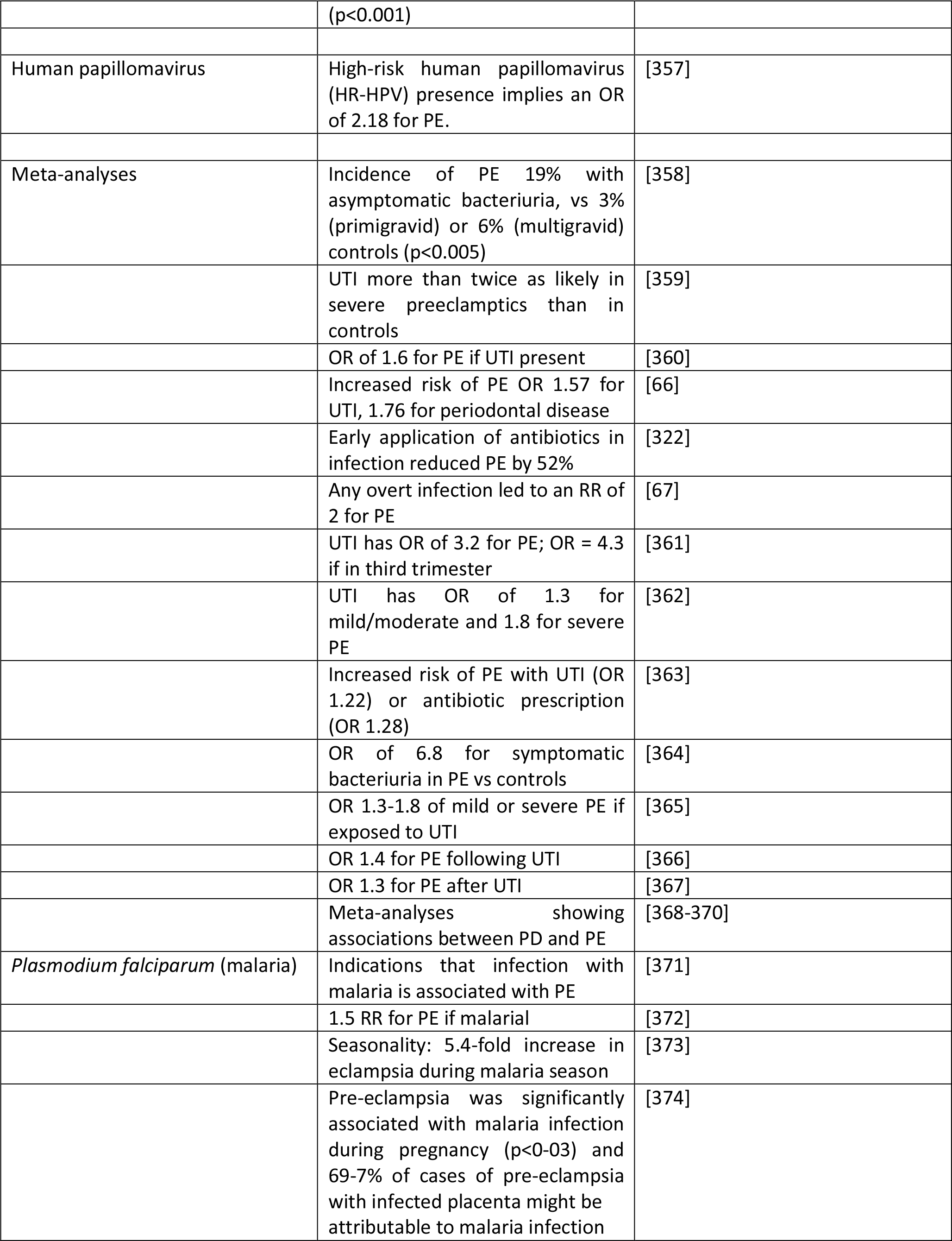

In contrast to the situation in PE, albeit severe PE is associated with iatrogenic pre-term births, there is a widespread recognition (e.g. [375–402]) that infection is a common precursor to pre-term birth (PTB) in the absence of PE. The failure of antibiotics to help can be ascribed to their difficulty of penetrating to the trophoblasts and placental regions. Unfortunately no proteomic biomarkers have yet been observed as predictive of PTB [403; 404]. In a similar vein, and if we are talking about a time of parturition that is very much more ‘preterm’, we are in the realm of miscarriages and spontaneous abortions and stillbirths, where infection again remains a major cause [405–408]. Here we note that <u>early or pre-emptive</u> antibiotic therapy <u>has</u> also proved of considerable value in improving outcomes after multiple spontaneous abortions [409].

## Vaginal, placental and amniotic fluid microbiomes in PE

It might be natural to assume that the placenta is a sterile organ, like blood is supposed to be. However, various studies have shown the presence of microbes in tissues including the placenta [386; 395; 410–422], vagina [383; 423–429], uterus [387; 430; 431], amniotic fluid [422; 432–437], and follicular fluid [438; 439], and how these may vary significantly in PE (we do not discuss other pregnancy disorders such as small for gestational age (SGA) and intrauterine growth restriction (IUGR)). We list some of these in Table 3.

**Table 3.**
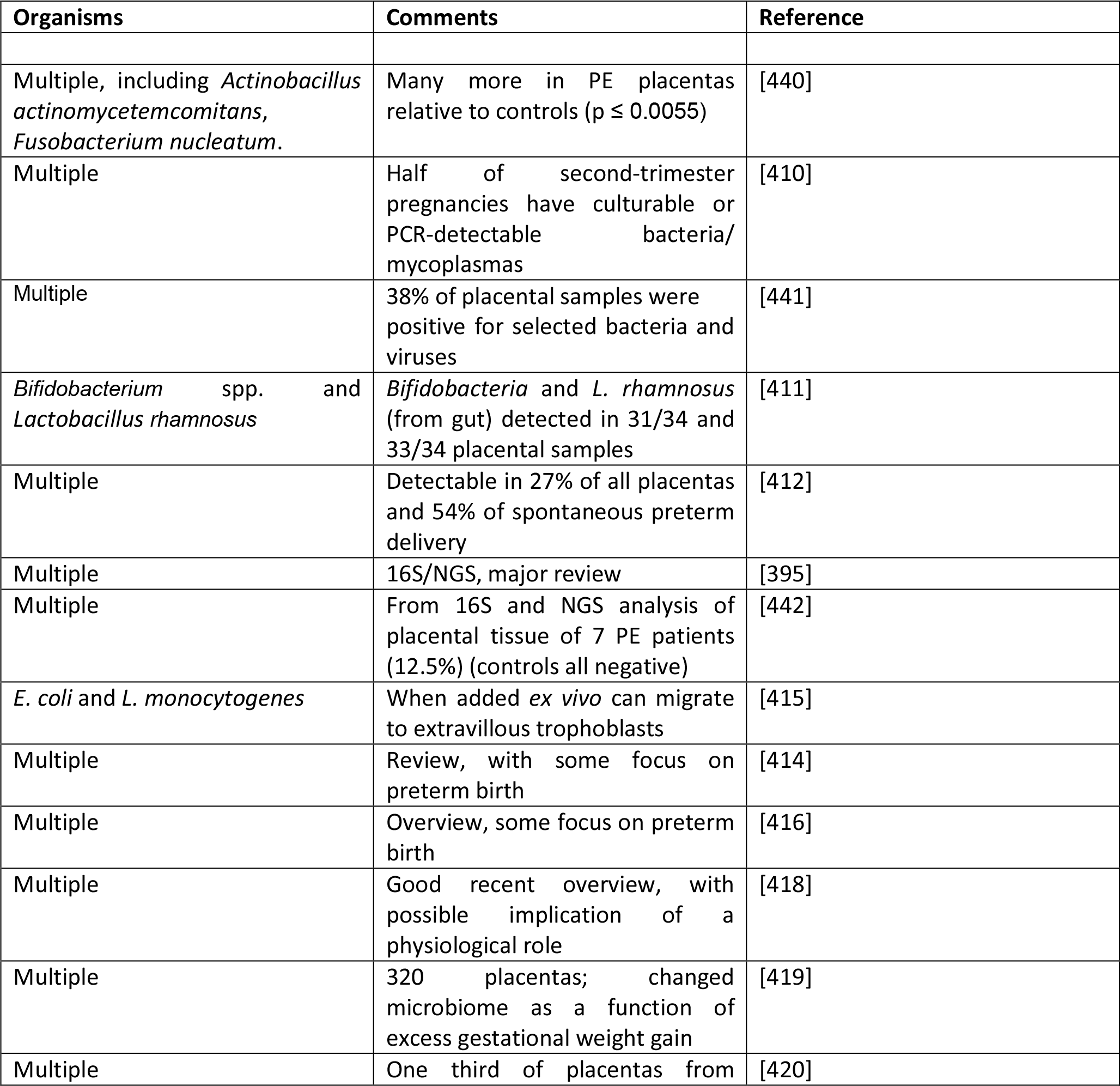
Evidence for microbes in placental tissues, including those with PE.

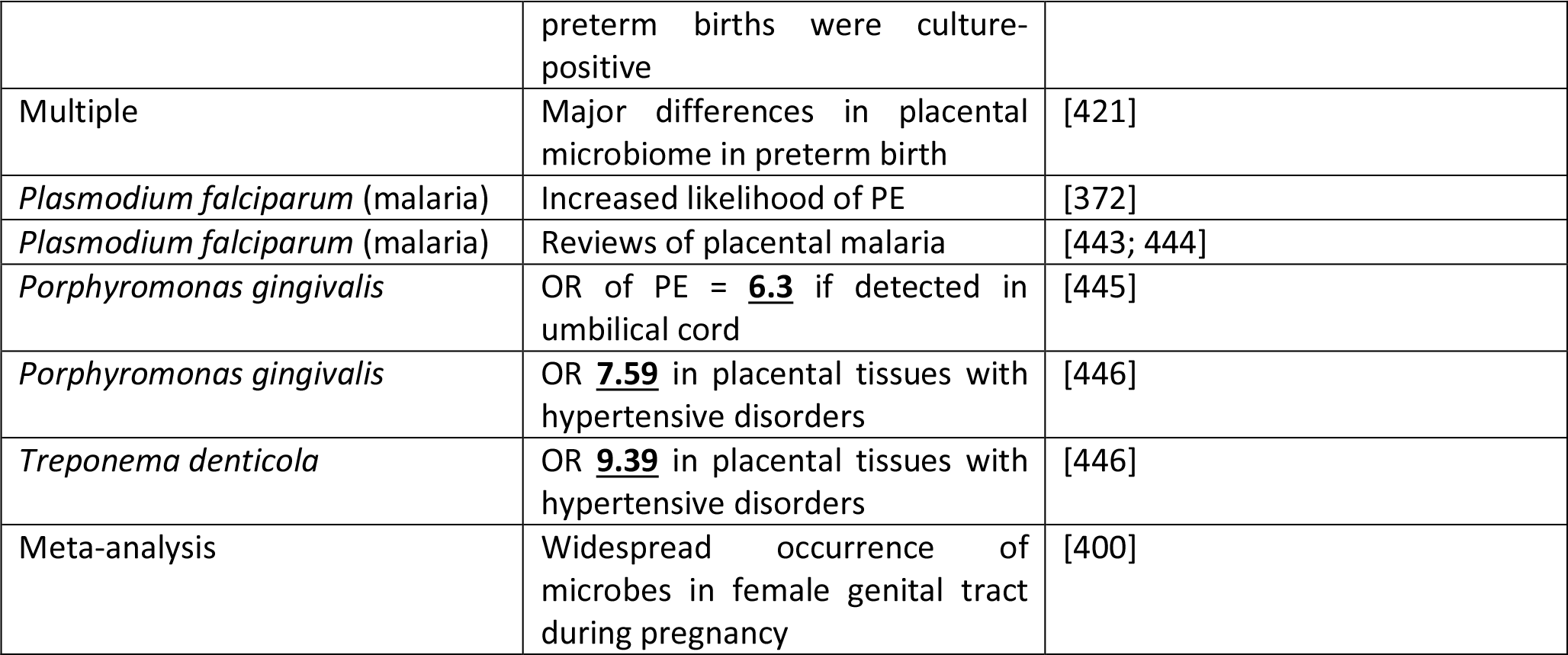

## Origins of a blood and tissue microbiome

As assessed previously [132–134] over a large literature, the chief source of blood microbes is the gut [418], with another major entry point being via the oral microbiome (especially in periodontitis, see below). For rheumatoid arthritis [135; 447–449] and diseases of pregnancy, urinary tract infection (see below and Table TT) also provides a major source.

## Gut origins of blood microbes and LPS

We have recently rehearsed these issues elsewhere [132–134], so a brief summary will suffice. Clearly the gut holds trillions of microbes, with many attendant varieties of LPS [450], so even low levels of translocation (e.g. [451–453]), typically via Peyer’s patches and M cells, provide a major source of the blood microbiome. This may be exacerbated by intra-abdominal hypertension that can indeed stimulate the translocation of LPS [454]. For reasons of space and scope, we do not discuss the origins and translocation of microbes in breast milk [455], nor the important question of the establishment of a well-functioning microbiome in the foetus and neonate [456], and the physiological role of the mother therein.

## Pre-eclampsia and periodontal disease

One potential origin of microbes that might be involved in, or represent a major cause of, pre-eclampsia is the oral cavity, and in particular when there is oral disease (such as periodontitis and gum bleeding) that can allow microbes to enter the bloodstream. If this is a regular occurrence one would predict that PE would be much more prevalent in patients with pre-existing periodontitis (but cf. [457] for those in pregnancy) than in matched controls; this is indeed the case (table 4).

**Table 4.**
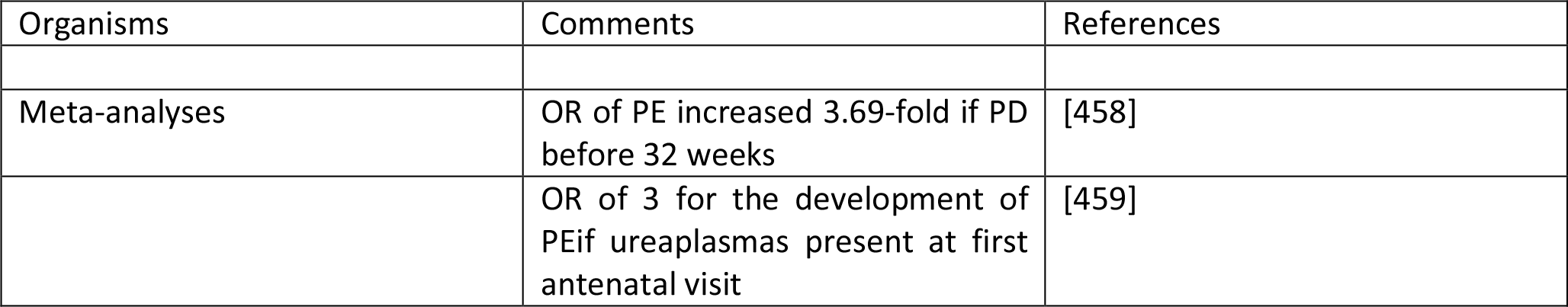
Periodontal disease (PD) and pre-eclampsiaUrinary tract infections (UTI)

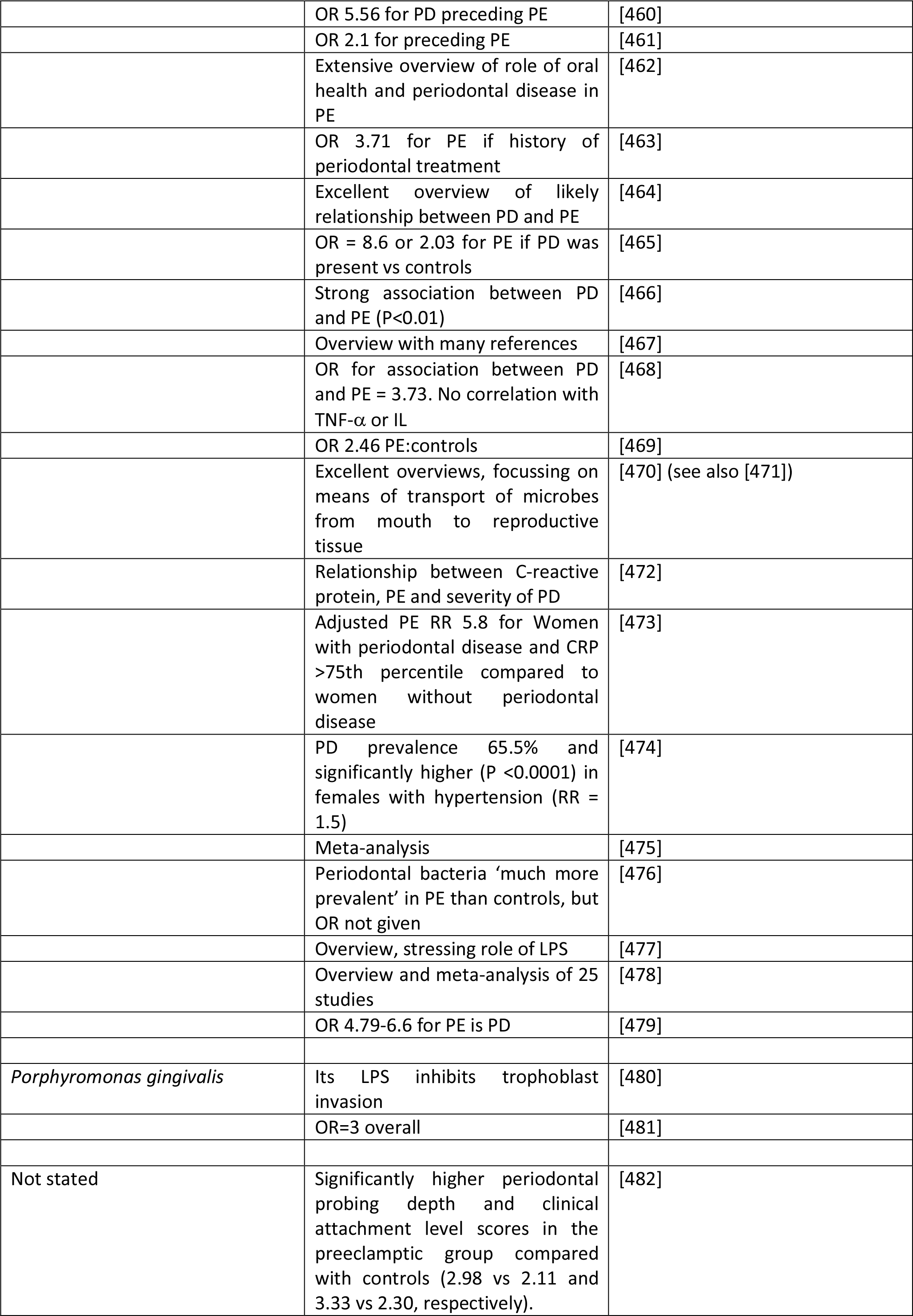

## Urinary tract infections (UTI)

A particular feature of UTIs is the frequency of reinfection [483–490]. This is because the organisms can effectively ‘hide’ in bladder epithelial cells as so-called ‘quiescent intracellular reservoirs’ [206; 487; 489; 491–495] of (presumably) dormant cells that can resuscitate. This is why reinfection is often from the same strains that caused the original infection [496–500]. Other complications can include renal scarring [501]. Bacteriuria (often asymptomatic) is a frequent occurrence in pregnancy (e.g. [365; 367; 459; 502–508]), and the frequency of UTI as a source of microbes causing PE is clear from Table 2.

## From blood to and from the placenta; a role for microparticles

We and others have noted the fact that many chronic, inflammatory disease are accompanied by the shedding of various antigens and other factors; typically they pass through the bloodstream as microparticles [119; 133; 509–514], sometimes known as endosomes [337; 339; 340; 510; 515] (and see later under miRNAs). Similarly, LPS is normally bound to proteins such as the LPS-binding protein and apoE (see [133]).

## Evidence from antibiotic therapies

Antibiotic drug prescriptions may be seen as a proxy for maternal infection, so if dormant (and resuscitating and growing) bacteria are a major part of PE aetiology one might imagine an association between antibiotic prescriptions and PE. According to an opposite argument, antibiotics and antibiotic prescriptions given for nominally unrelated infections (UTI, chest, etc, and in particular diseases requiring **<u>long-term</u>** anti-infective medication that might even last throughout a pregnancy) might have the beneficial side-effect of controlling the proliferation of dormant cells as they seek to resuscitate. There is indeed some good evidence for both of these, implying that it is necessary to look quite closely at the nature, timing and duration of the infections and of the anti-infective therapy relative to pregnancy. A summary is given in (Table 5. A confounding factor can be that some (e.g. the antiretroviral) therapies are themselves quite toxic [516; 517]; while the OR for avoiding PE was 15.3 in one study of untreated HIV-infected individuals vs controls, implying (as is known) a strong involvement of the immune system in PE, the ‘advantage’ virtually disappeared upon triple-antiretroviral therapy [518]. Overall, it is hard to draw conclusions from antiretrovirals [519; 520]. However, we have included one HIV study in the Table. Despite a detailed survey, we found no reliable studies with diseases such as Lyme disease or tuberculosis, where treatment regimes are lengthy, that allowed a fair conclusion as to whether antibiotic treatment was protective against PE. However, we do highlight the absolutely stand-out study of Todros and colleagues [521], who noted that extended spiramycin treatment (of patients with *Toxoplasma gondii*) gave a greater than tenfold protection against PE, when the parasite alone had no effect [522]. This makes such an endeavour (assessing the utility of early or pre-emptive antibiotics in PE) potentially highly worthwhile.

**Table 5.**
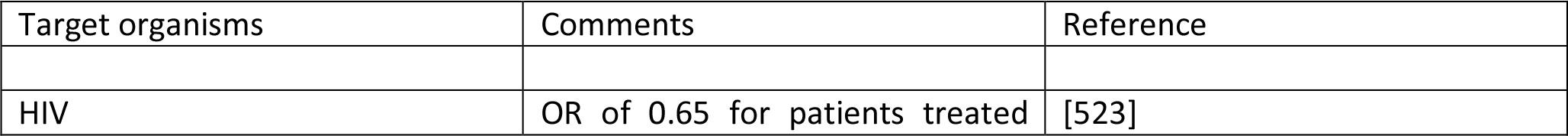
Examples of decreased PE following antibiotic therapies given for various reasons

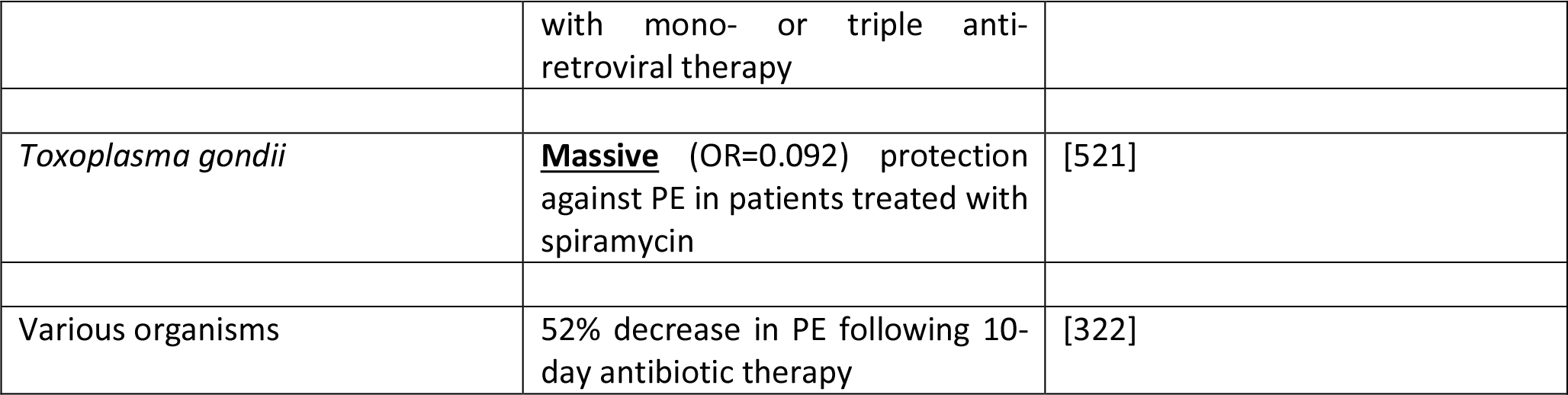

## Role of LPS in PE

It is exceptionally well known that LPS (*sensu lato*) is highly inflammagenic, and since one of us recently reviewed that literature *in extenso* [133] this is not directly rehearsed here. However, since we are arguing that it has a major role in PE naturally or *in vivo*, we do need to ask whether the literature is consistent with this more focussed question. The answer is, of course, a resounding ‘yes’. Notwithstanding that only primates, and really only humans, are afflicted by ‘genuine’ PE, so the genuine utility of rodent models is questionable [524], even if some can recapitulate elements of the disease [525; 526]. Hence, it is somewhat ironic that there are a number of animal models in which LPS (also known as ‘endotoxin’) is used experimentally to induce a condition resembling PE (e.g. [527–532], and see [533]). We merely argue that it is not a coincidence that exogenous administration of LPS has these effects, because we consider that it is in fact <u>normally</u> one of the main mediators of PE.

The standard sequelae of LPS activation, e.g. TLR signalling and cytokine production, also occur in PE [534; 535], bolstering the argument that this is precisely what is going on. In a similar vein, double stranded RNA-mediated activation of TLR3 and TLR7/8 can play a key role in the development of PE [536–538]. What is new here is our recognition that LPS and other inflammagens (e.g. [539–541]) may continue to be produced and shed by dormant and resuscitating bacteria that are generally invisible to classical microbiology.

## Effects of LPS and other microbial antigens on disrupting trophoblast invasion and/or stimulating parturition

As with other cases of cross-reactivity such as that of various antigens in *Proteus* spp that cause disease in rheumatoid arthritis [447–449], the assumption is that various microbial antigens can lead to the production of (auto-)antibodies that attack the host, in the present case of interest by stopping the placentation by trophoblasts. This is commonly referred to as ‘molecular mimicry’ (e.g. [542–545]), and may extend between molecular classes e.g. peptide/carbohydrate [546; 547]. Table 6 shows some molecular examples where this has been demonstrated.

**Table 6.**
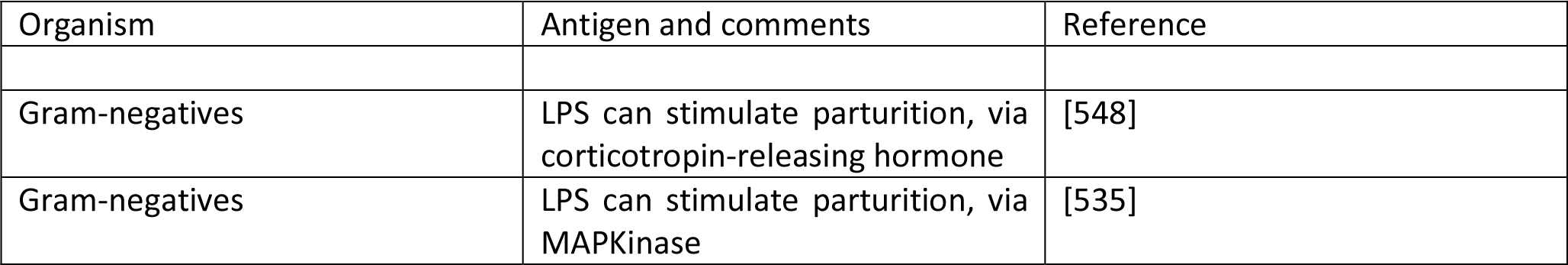
Molecular examples of bacterial antigens that can elicit antibodies that stop successful trophoblast implantation or stimulate parturition.

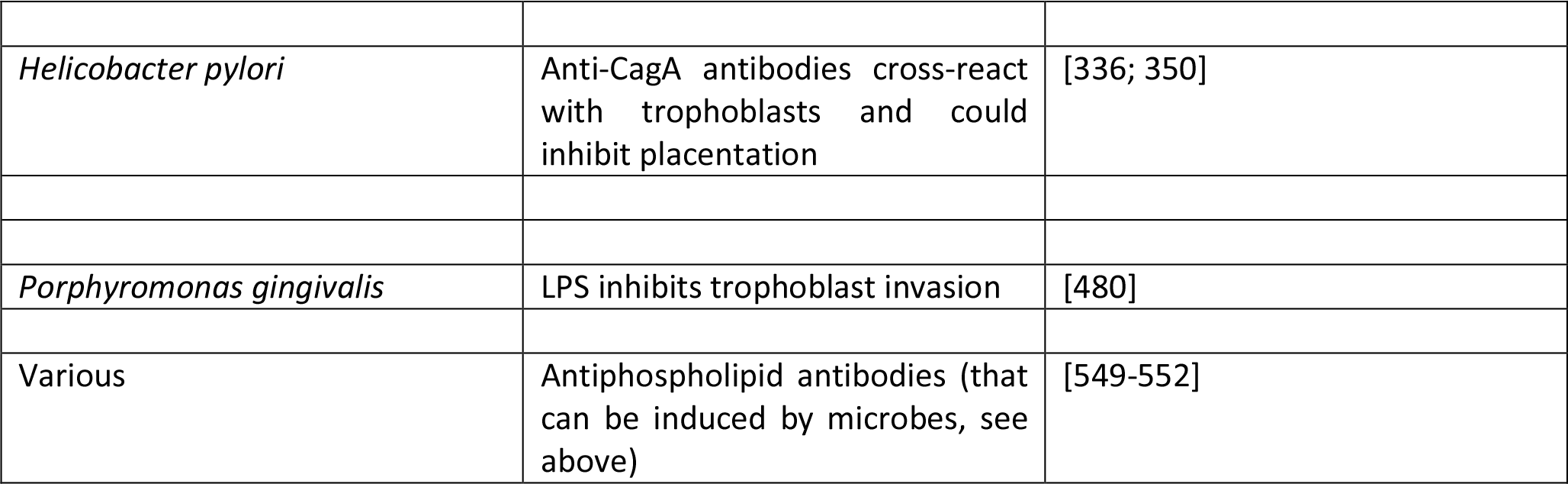

In many cases, the actual (and possibly microbial) antigens are unknown, and clearly the microbial elicitation of antibodies to anything that might contribute to PE points to multiple potential origins. To this end, we note that PE has also been associated with antibodies to angiotensin receptors [553–566], to smooth muscle [567; 568] (such blocking may be anti-inflammatory [569–571]), to adrenoceptors [572], to the M2 muscarinic receptor [573], and to Th17 [574](and see [575]). It is not unreasonable that epitope scanning of the antibody targets coupled to comparative sequence analysis of potential microbes might light up those responsible. In the case of Angiotensin II Type 1 Receptor Antibodies the epitope is considered [576] to be AFHYESQ, an epitope that also appears on parvovirus B19 capsid proteins; in the event, parvoviruses seem not to be the culprits here [577]. However, the role of these antibodies in activating the angiotensin receptor is also considered to underpin the lowering of the renin-angiotensin system that is commonly seen in PE [578–581], but which is typically raised during normal pregnancy.

Th-17 is of especial interest here, since these are the helper T (Th)-cell subset that produce IL-17. IL-17 is probably best known for its role in inflammation and automimmunity [575; 582–586]. However, it also has an important role in induction of the protective immune response against extracellular bacteria or fungal pathogens at mucosal surfaces [584; 587–599]. Th17 cells seem to participate in successful pregnancy processes and can be lower in PE [600–602], though more studies show them as higher [575; 603–611] or unchanged [612; 613]. One interpretation, consistent with the present thesis, is that the antimicrobial effects of <u>placental</u> IL-17 relative to T_reg_ cells are compromised during PE [575; 609; 614].

**A note on the terminology of sepsis**. As one may suppose from the name, sepsis (and the use of words like ‘antiseptic’) was originally taken to indicate the presence of culturable organisms in (or in a sample taken from) a host, e.g. as in bacteraemia. Recognising that it is the <u>products</u> of bacteria, especially cell wall components, that cause the cytokine storms that eventually lead to death from all kinds of infection [615–618], ‘sepsis’ nowadays has more come to indicate the latter, as a stage (in the case of established infection) on a road that leads to septic shock and (eventually) to death (with a shockingly high mortality, and many failures of initially promising treatments, e.g. [619; 620], and despite the clear utility of iron chelation [79; 118; 621; 622]). In most cases significant numbers of culturable microbes are either unmeasured or absent, and like most authors we shall use ‘sepsis’ to imply the results of an infection whether the organisms are detected or otherwise. Overall, it is possible to see the stages of PE as a milder form of the sepsis cascade on the left-hand side of Figure 7. Fig 7 compares the classical route of sepsis-induced death with the milder versions that we see in PE; they are at least consistent with the idea that PE is strongly related to the more classical sepsis in degree rather than in kind.

**Figure 7.**
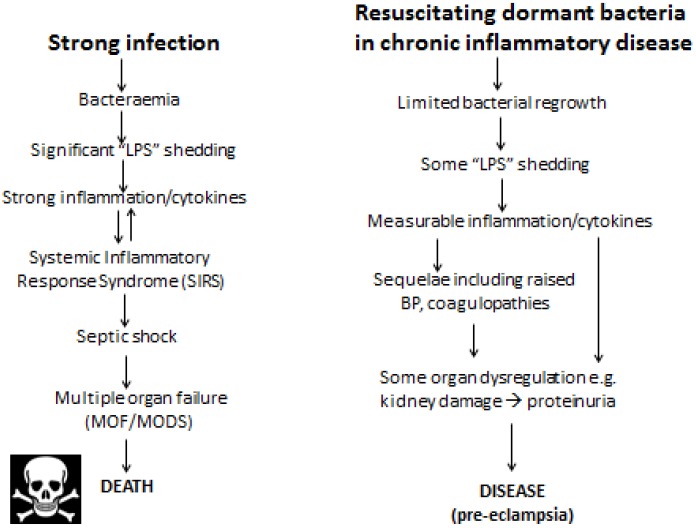
Pre-eclampsia bears some similarities to, and may be considered as a milder form of, the changes that occur during genuine sepsis leading to a systematic inflammatyory response syndrome, septic shock and multiple organ disfunction.

**Pre-eclampsia and neonatal sepsis**. If PE is really based on infectious agents, it is reasonable that one might expect to see a greater incidence of neonatal sepsis (i.e. infection) following PE. While there are clearly other possible explanations (e.g. simply a weakened immune system, sometimes expressed as neutropaenia, after PE), there is certainly evidence that this is consistent with this suggestion [623–627].

**PE biomarkers and infection**. Because of the lengthy development of PE during pregnancy, there has long been a search for biomarkers (somewhat equivalent to the ‘risk factors’ discussed earlier) that might have predictive power, and some of these, at both metabolome [14; 628–635] and proteome [636–638] level, are starting to come forward. The typical experimental design is a case-control, in which markers that are raised or lowered significantly relative to the age-matched controls are considered to be candidate markers of PE. However, just as noted with leukocyte markers [76] and PCOS [639], that does not mean that they might not also be markers for other things too, such as infection [640]!

Thus, one prediction is that if dormant and resuscitating bacteria are responsible for PE then **at least some of these biomarkers should also be (known to be) associated with infection**. However, one obvious point is that the markers may appear only <u>after</u> infection, and this may itself be after the first trimester; clearly then these would not then be seen as ‘first-trimester’ biomarkers! There are many well-known inflammatory biomarkers that are part of the innate (and possibly trained [641]) immune response, such as the inflammatory cytokines CRP (cf. [642; 643]), IL-6 [644], IL-1β [645], TNFα [646], and macrophage migration inhibitory factor (MIF) [647], that are also all biomarkers of infection [648–652]. Certainly the fact that these increase in PE is consistent with a role for an infectious component. However, we shall mainly look at <u>other</u> biomarkers that are known to increase with PE, and see if they are also known to be biomarkers for (or at least changed in the presence of) infection (and see Th17/IL-17 above), and we next examine this. We shall see that pretty well every biomarker that is changed significantly in PE is also changed following infection, a series of findings that we consider adds very strong weight to our arguments.

## Proteomic and similar biomarkers-circulating and placental

What is really needed is a full systems biology strategy (see e.g. [93; 653–655]) that brings together the actors that interact then parametrises the nature of those interactions in a suitable encoding (e.g. SBML [656]) that permits their modelling, at least as an ODE model using software such as CellDesigner [657], COPASI [658] or Cytoscape [659]. Thus, to take a small example, “agonistic autoantibodies against the angiotensin II type 1 receptor autoantibodies (AT1-AA) are described. They induce NADPH oxidase and the MAPK/ERK pathway leading to NF-κB and tissue factor activation. AT1-AA are detectable in animal models of PE and are responsible for elevation of soluble fms-related tyrosine kinase-1 (sFlt1) and soluble endoglin (sEng), oxidative stress, and endothelin-1, all of which are enhanced in pre-eclamptic women. AT1-AA can be detected in pregnancies with abnormal uterine perfusion” [565]. Many such players have been invoked, and we next list some.

***Activin A.*** Activin A is a member of the transforming growth factor (TGF)-β superfamily. Its levels are raised significantly in PE [112; 660]. However, activin A is also well-established as a biomarker of infection [661–664].

***Calretinin.*** In a proteomic study of pre-clamptic vs normal placentas [665], calretinin was one of the most differentially upregulated proteins (P = 1.6.10^−13^ for preterm PE vs controls, P = 8.9.10^−7^ for term PE vs controls), and in a manner that correlated with the severity of disease. While calretinin (normally more expressed in neural tissue and mesotheliomas [666]) is not normally seen as a marker of infection, it is in fact raised significantly when *Chlamydia pneumoniae* infects human mesothelial cells [667].

***Chemerin*** is a relatively recently discovered adipokine, whose level can increase dramatically in the first trimester of pre-eclamptic pregnancies [668], and beyond [669]. Its levels are related to the severity of the pre-eclampsia [670–672]. Specifically, an ROC curve [673] analysis showed that a serum chemerin level >183.5 ng.mL^−1^ predicted pre-eclampsia with 87.8% sensitivity and 75.7% specificity (AUC, 0.845; 95% CI, 0. 811-0.875) [668]. Papers showing that chemerin is also increased by infection (hence inflammation) include [674; 675]; it even has antibacterial properties [676; 677], and was protective in a skin model of infection [678; 679]. In a study of patients with sepsis [680], circulating chemerin was increased 1.69-fold compared with controls (p = 0.012), and was also protective as judged by survival. These seem like particularly potent argument for a role of chemerin as a marker of infection rather than of pre-eclampsia *per se*, and for the consequent fact that PE follows infection and not *vice versa.*

***Cystatin C***. Not least because kidney function is impaired in PE, low MW proteins may serve as biomarkers for it. To this end, cystatin C (13 kDa) has been found to be raised significantly in PE [681–687]; it also contributed to the marker set in the SCOPE study [7; 15]. Notably, although it certainly can be raised during infection [688], it seems to be more of a marker of inflammation or kidney function [689; 690].

***D-dimer***. “D-dimer” is a term used to describe quite varying forms of fibrin degradation product(s) [691]. Given that PE is accompanied by coagulopathies, it is probably not surprising that D-dimer levels are raised in PE [692–696], though this is true for many conditions [697], and some of the assays would bear improvement [698; 699]. Needless to say, however, raised D-dimer levels are also a strong marker for infection [700; 701].

***Endoglin***. Endoglin is the product of a gene implicated [702; 703] in the rare disease Hereditary H**<u>a</u>**emorrhagic Telangiectasia. The role of endoglin remains somewhat enigmatic [704]. However, endoglin levels were 2.5-fold higher in pre<u>-</u>eclamptic placentas compared to normal pregnancies (15.4 ± 2.6 versus 5.7 ± 1.0, p < 0.01). After the onset of clinical disease, the mean serum level of soluble endoglin in women with preterm PE was 46.4 ng.mL^−1^, as compared with 9.8 ng.mL^−1^ in controls (P<0.001) [83]. Women with a particular endoglin SNP (AA) were 2.29 times more likely to develop PE than those with the GG genotype (P = 0.008) [705], and endoglin is seen as a reasonably good marker for PE [83; 660; 706–709] (cf. [710]). Again, endoglin levels are raised following infection by a variety of organisms [711–714], with a particularly clear example that it is a marker of infection coming from the fact that there is raised endoglin only in infected vs aseptic loosening in joints following arthroplasty [715]. In general, it seems likely that these circulating (anti)angiogenic factors are more or less markers of endothelial cell damage, just as we have described for serum ferritin [119].

***Ferritin***. The natural iron transporter in blood is transferrin (e.g. [716–721]), present at ca 1-2g.L^−1^, with ferritin being an intracellular iron storage molecule, so one is led to wonder why there is even any serum ferritin at all [119; 722]. The answer is almost certainly that it is a leakage molecule from damaged cells [119], and when in serum it is found to have lost its iron content [723–726]. Serum ferritin is, as expected, raised during PE [237; 239; 242; 246; 248; 727; 728] and in many other inflammatory diseases [119], including infection (e.g. [729; 730] and above).

***miRNAs*** MicroRNAs are a relatively novel and highly important class of ^~^22nt noncoding, regulatory molecules [731–734]. Some are placenta-specific, and those in the circulation (often in endo/exosomes [735–737]) can be identified during pregnancy [738–741], potentially providing a minimally invasive readout of placental condition [742–744]. There is aberrant expression of placenta-specific microRNAs (miRNAs) in PE including miR-517a/b and miR-517c [745–751] and miR-1233 [752]. C19MC is one of the largest miRNA gene clusters in humans, maps to chromosome 19q13.41, and spans a ~100 kb long region. C19MC miRNAs are processed from the cluster [753], are primate-specific, conserved in humans, and comprise 46 miRNA genes, including the miR-517 family [754]. miR-517 is known to be antiviral [755; 756], while miR-517a overexpression is apoptotic [757] and can inhibit trophoblast invasion [758]. Importantly for our argument, miR-517 molecules are overexpressed following infection [759; 760].

**Neuropeptide Y**. Although, as its name suggests, neuropeptide Y is a neurotransmitter it is also correlated with stress. Certainly it is related to noradrenaline (see below) that may itself be responsible for the raised BP in PE [761]. It is also raised in sepsis, where it is considered to counterbalance the vasodilation characteristic of septic shock (e.g. [762; 763]). The apparent paradox of a raised BP in PE and a lowered one in septic shock is considered to be related to the very different concentrations of endotoxin involved (Fig 7).

***NGAL (lipocalin 2, siderocalin)***. NGAL (neutrophil gelatinase-associated lipocalin) is a lipocalin that is capable of binding catecholate-based siderophores (see [118; 764; 765]). As such it is anti-microbial, and is also an inflammatory or sepsis biomarker [766; 767]. Given our interest in iron, it is not surprising that it is changed during PE. While one study suggested it to be decreased in PE [768], a great many other studies showed it to be increased significantly in PE, and typically in a manner that correlated with PE severity [686; 769–777]. Pertinently to PE, it is also well established as an early biomarker of acute kidney injury (AKI) [778–781]. However, it is not a <u>specific</u> biomarker for AKI vs sepsis [779; 782–790] and its origin in sepsis differs [791; 792]. Of course it can be the sepsis that leads to the AKI [793; 794]. Fairly obviously, while it does tend to be increased during PE, we again see its direct role as an antimicrobial and marker of sepsis as highly supportive of our present thesis.

***Placental growth factor (PlGF)***. This is a member of the vascular endothelial growth factor (VEGF) Family, that despite its name has a great many activities [795]. It is often considered in parallel with endoglin and sFlt, with a high sFlt:PlGF ratio being considered as especially discriminatory for PE [796–807], i. e. a lower PlGF can be diagnostic of PE [710; 808–810]. PlGF tends to be raised in sepsis unrelated to pregnancy [811; 812], while its lowering in PE may be due to the excess sFLT that decreases it [795; 813; 814]. In one study of a patient with CMV infection and PE it was in fact raised [815], while in the case of IUGR it was massively lowered [816]. PlGF alone is thus probably not a useful general marker for either PE or sepsis if one is trying to disentangle them, although it has clear promise when PE is superimposed on CKD [810; 817].

***Procalcitonin***. Procalcitonin is the 116 amino acid polypeptide precursor of calcitonin, a calcium regulatory hormone. It is another marker that has been observed to be raised (according to severity) in pre-eclamptics [693; 818; 819](but cf. [820]). However, it too is a known marker of bacterial infections or sepsis [818; 821–829].

***Serum amyloid A***. This is an inflammatory biomarker, that was shown to increase fourfold in PE in one study [830], was significantly raised in another [819], but not in a third [831]. However, it is a well-established (and potent) biomarker for infection/sepsis (e.g. [832–845]). Defective amyloid processing may be a hallmark of PE more generally [846], and of course amyloid can be induced by various microbes [309; 311; 847; 848] and their products [250].

***Soluble fms-like tyrosine kinase-1 (sFlt)***. The soluble fms-like tryrosine kinase (sFlt) receptor is a splice variant of the VEGF receptor [706]. It is raised considerably in PE [660; 708; 796; 801; 803; 849–852], and may be causal [525; 566; 853–856]. Needless to say, by now, we can see that it is also a very clear marker of infection [708; 857; 858], whose levels even correlate with the severity of sepsis [859–861]. Of particular note is the fact that sFLT is actually <u>anti</u>-inflammatory [860].

***Thrombomodulin***. Soluble thrombomodulin was recognised early as an endothelial damage biomarker, and is raised in PE [862–872]. Interestingly, it has been found to have significant efficacy in the treatment of sepsis(-based DIC) [873–881].

***TLR4*** upregulation in preeclamptic placentas [882] is entirely consistent with infection and the ‘danger model’ as applied to PE [883]. As well as LPS activation (reviewed in [133]), the heat shock protein 60 of *Chlamydia* also activates TLR4 [131].

***Visfatin*** is another adipokine that is raised in PE, approximately two-fold in the study of Fasshauer and colleagues [884], and 1.5-fold in that of Adali and colleagues [885]. However, it was little different in a third study [886] while in a different study it was rather lower in pre-eclampsia than in controls [887]. This kind of phenomenon rather lights up the need for excellent quality studies, including ELISA reagents, when making assessments of this type.

Fairly obviously, the conclusion that this long list of biomarkers that are raised in PE might be <u>specific</u> ‘PE’ biomarkers is challenged very strongly by the finding that they are, in fact, all known markers of infection, a finding that in our view strongly bolsters the case for an infectious component in PE.

In a similar vein, there are a number of other sepsis markers (where sepsis is varied via or occurs as an independent variable) that we would predict are likely to be visible as raised in PE patient. These might include [652; 888] PAI-1, sE-selectin [889] and sVCAM-1 [859]. In particular, Presepsin looks like a potentially useful marker for sepsis [826; 827; 890–895] but we can find no literature on its use as a PE biomarker, where we predict that it may also be raised.

## Metabolomic biomarkers

For fundamental reasons connected with metabolic control and its formal, mathematical analysis [896–899], changes in the metabolome are both expected [900] and found [901–904] to be amplified relative to those in the transcriptome and proteome. For similar reasons, and coupled to evolution’s selection for robustness [905–911] (i.e. homeostasis) in metabolic networks, we do not normally expect to find single metabolic biomarkers for a complex disease or syndrome. Since our initial metabolomic analyses [628], the technology has improved considerably [912–915], a full human metabolic network reconstruction has been published [911; 916–918] in the style of that done for yeast [919], and a number of candidate metabolomics biomarkers for PE have been identified reproducibly on an entirely separate validation set [14; 629].

This latter, LC-MS-based, study [14] found a cohort of 14 metabolites from the first trimester that when combined gave an OR of 23 as being predictive of third trimester PE. For convenience, we list them in Table 7. Note that because they were characterised solely via their mass there are some uncertainties in the exact identification in some cases, and that untargeted metabolomics of this type has a moderately high limit of detection (maybe 10 μM) such that many potentially discriminatory metabolites are below the limit of detection.

**Table 7.**
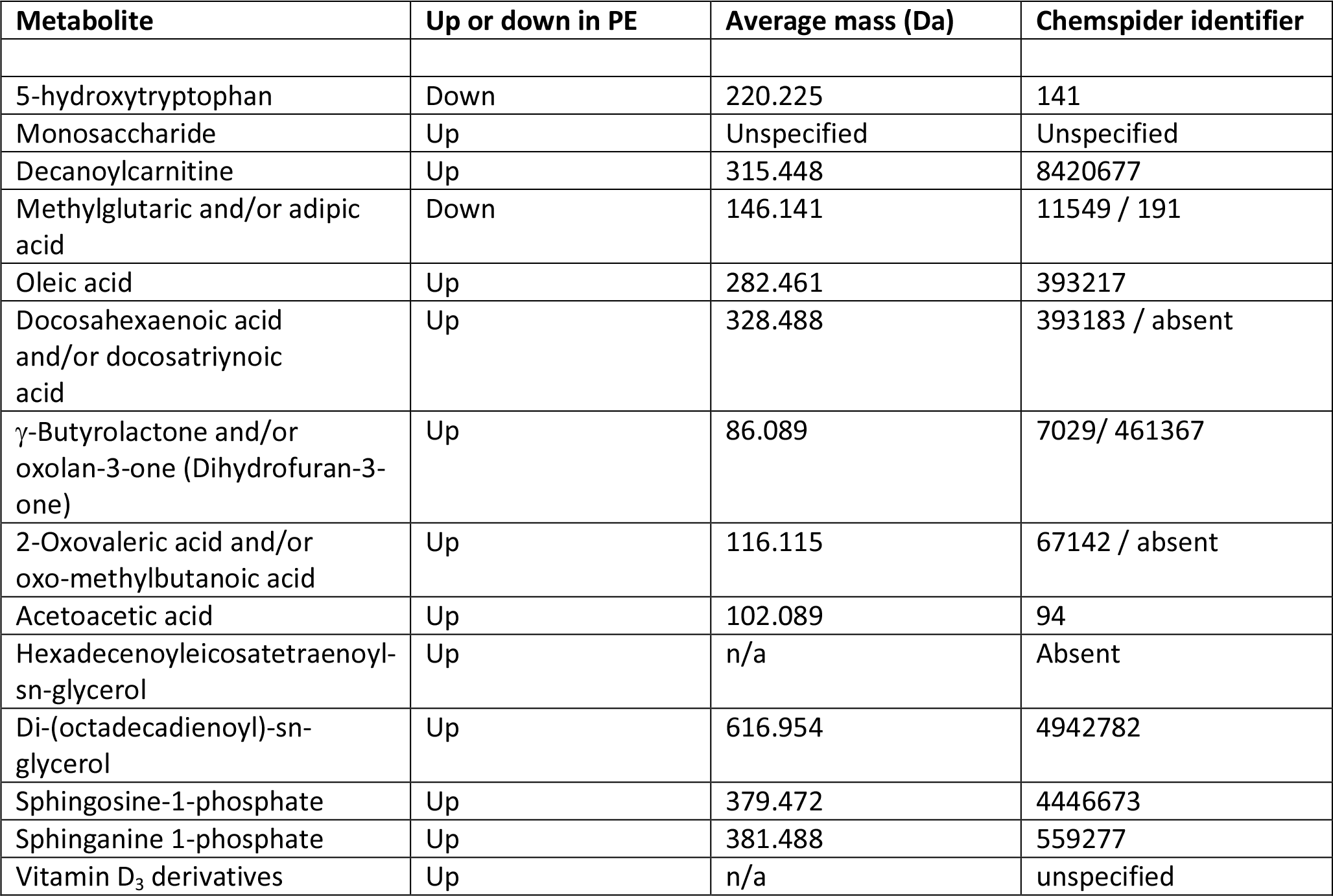
14 metabolites contributing to a pre-eclamptic ‘signature’ [14]

A number of features of interest emerge from this.

1. All the markers save 5-hydroxytryptophan and adipic/methylglutaric acid were raised in PE; 5-hydroxytryptophan is a precursor of serotonin (which in some studies [920] has been seen to be mildly elevated in PE).
2. Markers came from multiple classes of metabolite or areas of metabolism, including amino acids, carbohydrates, carnitines, dicarboxylic acids, fatty acids (especially), (phospho)lipids and sterols.
3. γ-Butyrolactone derivatives can act as signalling molecules for a variety of bacteria [921; 922].
4. In common with many other inflammatory diseases [138], Vitamin D_3_ levels (usually measured as 25(OH)vitD or calcidiol) are often lower in PE [923–927](cf. [928–930]), consistent with the levels of their derivatives being raised. However, the direction of causality inflammation **←→** vitamin D levels is not yet known [931] (see also [136; 138; 930]).
5. None of these metabolites was among four metabolites proposed as first trimester biomarkers in two other (smaller) studies from different groups [634; 932].
6. Sphingolipid metabolism can be deranged in PE [933] (also in Parkinson’s [934]).

As well as the non-targeted metabolomics noted above, a number of other small molecule biomarkers have been turned up by more conventional measurements.

***Noradrenaline (norepinephrine)***. An interesting early study [935] found that venous plasma noradrenaline was raised by 67% in pre-eclamptics vs controls. Similar data were found by others [936]. This is of particular interest in the present context since noradrenaline is well established as highly growth stimulatory to Gram-negative microorganisms (e.g. [937–941]), in part by acting as a siderophore [942–944]. It also raises the levels of neuropeptide Y [761], and as a stress hormone [945], is of course well known for its role in raising blood pressure, a hallmark of PE.

There is relatively little metabolomics work in sepsis, but in one study carnitine and sphingolipid metabolism were also modified during sepsis [946], while in another [947] a suite of molecules were decreased during acute sepsis. However, the patients involved here were quite close to death, so it is not clear that comparisons between the metabolome in PE and in dying patients are that worthwhile.

We also note a recent and rather interesting suggestion by Eggers [948] that the maternal release of adrenaline (rather than noradrenaline) may have an important aetiological role in PE, although as with the rest of our thesis here it is not there indicated as to what causes the adrenaline to rise (although infection and inflammation can of course do so).

***Uric acid***. Hyperuricemia is a moderately common finding in preeclamptic pregnancies, and may even be involved in its pathogenesis (see e.g. [949–955]). However, it does not seem to be very specific [956–960], and is seemingly not an early biomarker (and it did not appear in our own study [14]). Its lack of specificity is illustrated by the fact that there is considerable evidence for the roles of purinergic signalling [961], and especially the role of uric acid, in Alzheimer’s and Parkinson’s disease [962–964], as well as in a variety of other kinds of inflammatory processes, including pro-inflammatory cytokine production [965; 966], the *Plasmodium falciparum-induced* inflammatory response [967], the mechanistic basis for the action of alum as an adjuvant [968], and even peanut allergy [969–971]. As is common in case-control studies when just one disease (e.g. PE) is studied, artificially high levels of sensitivity and (especially) specificity may appear when other patients with other diseases are not considered.

## Clotting, coagulopathies and fibrinogen in PE

In much of our previous work (e.g. [119–127]), we have noted that each of these chronic, inflammatory diseases is accompanied by changes in fibrin fibre morphologies, coagulopathies and changes in erythrocytes that are both substantial and characteristic. They can variously be mimicked by adding unliganded iron or LPS. As is well known, LPS itself is a strong inducer of coagulation, whether via tissue factor or otherwise (e.g. [972–981]), and will bind to fibrin strongly [252; 982]. The morphological methods have not yet to our knowledge been performed on blood from pre-eclamptics, whether as a diagnostic or a prognostic, though we note that clotting factors came top in one GWAS looking for gene-PE associations [111]. Fibrinogen itself is a TLR4 ligand [983], is raised in PE [984–987], and we note the extensive evidence for coagulopathies during pregnancies with PE (e.g. [25; 121; 511; 692; 988–1000]). In the worst cases these are the very frightening Disseminated Intravascular Coagulation (DIC) [980; 1001–1005], that can, of course, also emerge as a consequence of sepsis [1006–1012]. Variations in the plasminogen activator inhibitor-1 may contribute to the hypofibrinolysis observed [1013–1015].

We recently showed that LPS can potently induce amyloid formation in fibrin (see [251; 1016]). Thus, In addition, we note the increasing recognition that amyloid proteins themselves, that may occur as a result of coagulaopathies, are themselves both inflammatory (e.g. [540; 640; 1017–1022]) and cytotoxic (e.g. [250; 1023–1027]), and this that can of itself contribute strongly to the death of e.g. trophoblasts.

Related to clotting parameters are three other ‘old’ but easily measured variables that probably reflect inflammation [1028] and that have been suggested to differ in PE from normotensives and may have some predictive power. The first two are the erythrocyte sedimentation rate (ESR) [1029; 1030] and the red cell distribution width (RDW) [1031] (but cf. [1032]). Interestingly, the former was the only variable that was predictive of a subsequent stroke following sub-arachnoid haemorrhage [1033]. The third relates to the morphology of erythrocytes (that may in part underpin the other two). We and others have shown in a series of studies (e.g. [127–129; 1034–1037] that erythrocyte morphology diverges very considerably from that ‘classical’ discoid shape adopted by normal healthy cells, and that this can be a strong indicator of disease [130]. In extreme cases (e.g. [126; 1038–1043]), including following infection [1044], this results in eryptosis, the suicidal death of erythrocytes. It is of interest that ceramide, a precursor of sphingosine-1-phosphate (Table 7), is raised in various diseases such as Parkinson’s and may serve to stimulate eryptosis [1045]. Although we know of no direct measurements to date, there is evidence that eryptosis may play a significant role in PE [1046]

**Prevention Strategies**. Apart from low-dose aspirin (that may have little effect [1047–1050] unless initiated relatively early in pregnancy [1051–1055]), and low-dose calcium [1056], there are relatively few treatment options in present use [1057–1060]. (Magnesium sulphate [1061–1063] has been used as a treatment for eclampsia.)

In the history of science or medicine, some treatments are empirical, while others are considered to have a mechanistic basis. The general assumption is that the more we know about the originating aetiology of a disease or syndrome the more likely we are to be able to treat its causes effectively, and not just its symptoms. Clearly, too, clinicians are rightly loth to give complex and potentially teratogenic treatments to pregnant women when this can be avoided [1064; 1065]. However, the surprising lack of systematic data with antibiotics [1066], modulo one particularly spectacular success [521], suggests that we ought to be performing trials with safe antibiotics on women at special risk [1067]. These must take care to avoid any Jarisch-Herxheimer reaction [1068–1070] due to the release from microbes induced by antibiotics of inflammagens like LPS [1071–1074]. A related strategy recognises that some FDA-approved drugs can actually exert powerful antibiotic effects *in vivo* (but not on petri plates) <u>by modifying the host</u> [1075].

Because of the known oxidative stress accompanying PE, it had been assumed that antioxidants such as vitamin C (ascorbate) might be preventive; however, this turned out not to be the case (even the opposite) for ascorbate [1047; 1076]. Probably this is because in the presence of unliganded iron, ascorbate is in fact pro-oxidant [118]. However, polyphenolic antioxidants that actually act by chelating iron [79; 118] seem to be more effective [1077].

Another area that we and others have previously highlighted recognises the ability of non-siderophoric iron chelators to act as iron-withholding agents and thereby limit the growth of bacteria. Again, a prediction is that women with iron overload diseases should be more susceptible to pre-eclampsia, a prediction that is borne out for α-thalassaemia [1078; 1079] though not apparently for hereditary haemochromatosis [1080]. However, the extent of use of chelators and degree of control of free iron thereby obtained is rarely recorded in any detail, so in truth it is difficult to draw conclusions.

The general benefits of nutritional iron chelators such as blueberries and other fruits and vegetables containing anthocyanins have been discussed elsewhere (e.g. [79; 118; 1081]).

How significant coagulopathies are to the aetiology of PE development (as opposed to providing merely an accompaniment) is not entirely clear, but on the basis that they are then anticoagulants would potentially assist, just as thrombomodulin does in DIC accompanying sepsis [879; 881; 1008; 1012]. Of course one of many effects of low-dose aspirin is to act as an anticoagulant. There is also evidence for the efficacy of heparin [5; 1058; 1082–1087], which is especially interesting given our highlighting of the role of coagulopathies in PE. Those anticoagulants that avoid bleeding [1088] are obviously of particular interest, while anything stopping the fibrin forming β-amyloid [251; 252] should serve as an especially useful antiinflammatory anticoagulant.

With a change in focus from function-first to target-first-based drug discovery [909], there has been an assumption that because a drug is (i) found to bind potently to a molecular target and (ii) has efficacy at a physiological level *in vivo*, the first process is thus responsible for the second. This has precisely no basis in logic (it is a logical fault known variously as “affirming the consequent” or “post hoc ergo propter hoc” [1089]). This is because the drug might be acting physiologically by any other means, since drug binding to proteins is typically quite promiscuous (e.g. [1090–1094]). Indeed, the average <u>known</u> number of binding sites for marketed drugs is six [201; 1095]. In particular, it is likely, from a network or systems pharmacology perspective (e.g. [911; 1096–1099], that successful drugs (like aspirin) are successful <u>precisely</u> because they hit multiple targets. The so-called‘statins’ provide a particularly good case in point [118].

It had long been known that the enzyme HMGCoA reductase exerted strong control on the biosynthetic flux to cholesterol, and that inhibiting it might lower the flux and steady-state cholesterol levels (as indeed it does). Notwithstanding that cholesterol alone is a poor predictor of cardiovascular disease [1100–1102]), especially in the normal range, HMGCoA reductase inhibitors have benefits in terms of decreasing the adverse events of various types of cardiovascular disease [1103]. Following an original discovery of natural products such as compactin (mevastatin) and lovastatin containing a group related to hydroxymethylglutaric acid (rather than a CoA version) that inhibited the enzyme [1104]), many variants with this (hydroxyl)methylglutaric substructure came to be produced, with the much larger‘rest’ of the molecule being considerably divergent (see Fig 8, where the MW values vary from 390.5 (mevastatin) to 558.6 (atorvastatin)). Despite this wide structural diversity (Fig 8) they are still collectively known as ‘statins’, and despite the wildly illogical assumption that they might all work in the same way(s). The fact that different statins can cause a variety of distinct expression profiles [1105] is anyway utterly inconsistent with a unitary mode of action. In particular, in this latter study, statins clustered into whether they were (fluvastatin, lovastatin and simvastatin) or were not (atorvastatin, pravastatin and rosuvastatin) likely to induce the side effect of rhabdomyolysis or any other myopathy. Clearly, any choice of ‘statin’ should come from the latter group, with pravastatin and rosuvastatin being comparatively hydrophilic.

**Figure 8.**
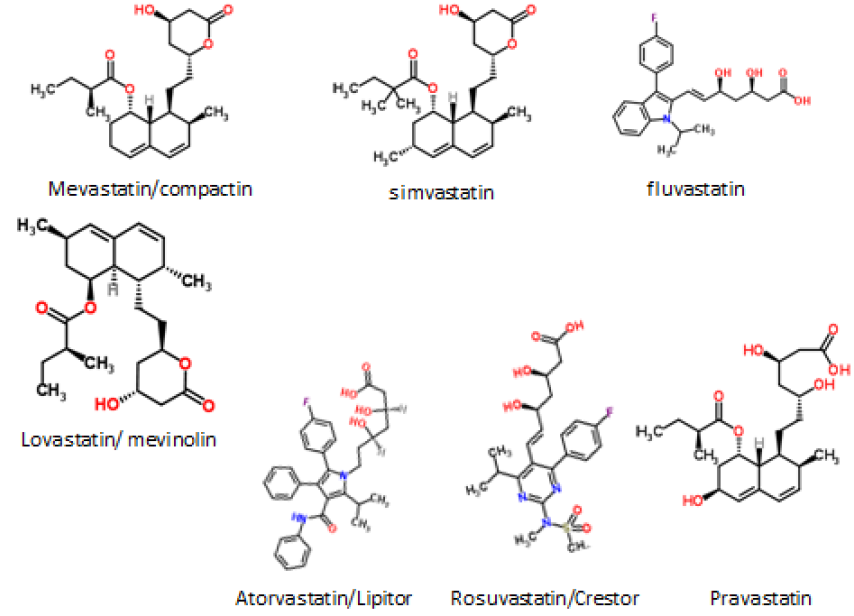
Some structures of various statins

The epidemiological fact of improved survival despite the comparative irrelevance of cholesterol levels to atherosclerotic plaque formation and heart disease in the normal range provides an apparent paradox [1106]. This is easily solved by the recognition (e.g. [1107–1120], and many other references and reviews) that ‘statins’ are in fact anti-inflammatory. They may also be antimicrobial/anti-septic, whether directly or otherwise [1121–1125], and we also note the role of cholesterol in mopping up endotoxin [1126]. Finally, here, it needs to be recognised that statins do themselves serve to lower iron levels [1127–1129], and (while oddly this seems not to have been tested directly) simple inspection of their structures (Fig 8) implies that the better ones (with their multiple OH groups) might in fact chelate iron directly.

In consequence, a number of authors have indicated the potential utility of statins in treating PE [107; 525; 1130–1140], and pravastatin has been the subject of a number of favourable studies [525; 1131; 1133; 1136; 1138; 1141; 1142], including in humans [1131; 1143–1145]. Pravastatin seems more than ripe for a proper, randomised clinical trial [1130].

Another ‘vascular’ class of drugs that has been proposed for treating PE is represented by those of the family of vasodilatory phosphodiesterase5 inhibitors such as sildenafil (Viagra) and vardenafil (Levitra), as it is reasonable that they might improve endothelial function, especially if started early in pregnancy [1146]. Thus vardefanil restores endothelial function by increasing placental growth factor [1147], and sildenafil has shown promise in a number of animal studies [1148–1153] and in human tissues [1154; 1155], with a clinical trial ongoing [1156]. In particular [1153], it was able to normalise the metabolomics changes observed in a mouse model (the COMT^-/-^ model) of PE.

Antihypertensive therapy for PE has been reviewed by Abalos and colleagues [1157] and Magee and colleagues [108]. Anti-hypertensives did halve the incidence of hypertension but had no effect on PE. Methyldopa is one of the most commonly used anti-hypertensives in pregnancy, but it may also stimulate eryptosis [1158]; alternative drugs were considered to be better [1157] for hypertension. Nifedipine [1159] and labetalol [1160] are considered a reasonable choice. There was also a slight reduction in the overall risk of developing proteinuria/pre-eclampsia when beta blockers and calcium channel blockers considered together (but not alone) were compared with methyldopa [1157]. In mice, olmesartan (together with captopril) proved usefully anti-hypertensive [1161]; this is of interest because olmesartan is also an agonist of the vitamin D receptor [1162]. However, it was not mentioned in either [1157] or [108].

LPS itself has long been recognised as a target of inflammatory diseases. Unfortunately, despite initially promising trials of an anti-LPS antibody known as Centoxin [1163], it was eventually withdrawn, apparently because of a combination of ineffectiveness [1164; 1165] and toxicity [1166; 1167]. LPS is rather hydrophobic, and thus it is hard to make even monoclonal antibodies very selective for such targets, such that the toxicity was probably because of its lack of specificity between lipid A and other hydrophobic ligands [1168]. Other possible treatments based on LPS, such as ‘sushi peptides’ [1169–1176] (or variants [1177; 1178]) and LPS-binding protein were covered elsewhere [133].

If an aberrant or dysbiotic gut microbiome is the source of the microbes that underpin PE, it is at least plausible that the gut microbiome should be predictive of PE [370], but we know of no suitably powered study that has been done to assess this, and this would clearly be worthwhile. However, in a study of primiparous women, the OR for getting severe PE was only 0.6 if probiotic milk drinks containing lactobacilli were consumed daily [1179]. This too seems an area well worth following up.

From a metabolomics point of view, the molecules seen to be raised in PE may either be biomarkers of the disease aetiology <u>or</u> of the body’s attempts to respond to the disease (and this is true generally [1180]). Thus it is of great interest that sphingosine-1-phosphate (S1P) was raised in PE (see [14] and Table 1). S1P is mainly vasoconstrictive [1181; 1182], but agonists of the sphingosine-1-phosphate1 receptor (that is involved in endothelial cell function) seemed to have considerable value in combatting the cytokine storm that followed infection-driven sepsis [1183–1188]. The detailed mechanism seems not to be known, but in the context of infection, a need for S1P and other sphingolipids for successful pregnancies [1189; 1190] (see also Parkinson’s [934]), and the induction of PE by its disruption [933; 1191–1195]), some serious investigation of the potential protective effects of S1PR1 agonists seems highly warranted.

Among other small molecules, melatonin has shown some promise in the treatment of septic shock, by lowering inflammatory cytokine production [1196] (and see [1197] for neonatal oxidative stress), and a trial is in prospect for PE [1198].

Lipoxin A_4_ (LXA_4_) is considered to be an endogenous stop signal in inflammation. While recognising the difficulties with rodent PE models (above), we note that in one study, the effect of BML-111 (a synthetic analogue of LXA_4_) was tested on experimental PE induced in rats by low-dose endotoxin (LPS), and showed highly beneficial effects [530].

**Coda-a return to the Bradford Hill criteria**. Returning to the Bradford Hill criteria for ascribing causation of a disease to an environmental factor [91], we can now ask whether a detectable (if largely dormant) microbiome X, that is more likely to replicate with free iron, and can anyway secrete or shed a variety of inflammatory components such as LPS, represents a plausible and major aetiological factor for PE (Y):

1. what is the strength of association between X and Y? We found an overwhelming co-occurrence of microbes or their products and PE
2. what is the consistency of association between X and Y? Almost wherever we looked, whether via periodontal disease, urinary tract infection, or other means of ingress, we could find a microbial component in PE
3. What is the specificity of association between X and Y? Insufficient data are available to ascribe PE solely to one type of organism; by contrast the data rather indicate that a variety of microbes, each capable of shedding inflammatory molecules such as LPS, can serve to stimulate or exacerbate PE.
4. experiments verify the relationship between X and Y. It is unethical to do these in humans in terms of purposely infecting pregnant women, but data from antibiotics show the expected improvements.
5. modification of X alters the occurrence of Y; this is really as (4)
6. biologically plausible cause and effect relationship. Yes, this is where we think the ideas set down here are entirely consistent with current thinking on the main causes of PE. What we add in particular is the recognition that bacteria (and other microbes) that may be invisible to culture are both present and responsible, by established means, for the inflammation and other sequelae (and especially the coagulopathies) seen as causative accompaniments to PE.

## Other predictions

Classical clinical microbiology, involving mainly replication-based methods, is evolving rapidly to assess the microbial content of samples on the basis of DNA sequences [288; 1199], including 16S rDNA [279; 280; 282; 284; 285; 287; 289; 1200], suitable protein-encoding housekeeping genes (e.g. [1201–1206]), and, increasingly, full genome sequences [1207]. In the future, we can thus expect a considerable increase in molecular assessments of the microbiological content of blood, urine and tissues, and this will obviously be a vital part of the experimental assessment and development of the ideas presented here. Molecular methods will also be used to assess maternal circulating DNA [1208–1210] and RNA [1211] in terms of both its presence and sequencing, as well as the use of digital PCR [1212].

Since PE has such a strong vascular component, we also predict that measurements designed to detect coagulopathies will increase in importance, for both diagnosis and prognosis, and for assessing treatments.

New drugs designed to kill <u>non-growing</u> bacteria [1213–1217] or to overcome amyloid coagulopathies [1218–1222] will be needed, and will come to the fore.

Finally, we consider that real progress in understanding PE from a systems biology perspective means that it must be modelled accordingly, and this must be a major goal.

**Concluding remarks**. We have brought together a large and widely dispersed literature to make the case that an important aetiological role in pre-eclampsia is played by dormant microbes, or at least ones that are somewhat refractory to culture, and that these can awaken, shed inflammagens such as LPS, and thereby initiate inflammatory cascades. (The sequelae of these, involving cytokines, coagulopathies, and so on, are well enough accepted.) The case is founded on a large substructure of interlocking evidence, but readers might find the following elements as discussed above especially persuasive and/or worthy of follow-up:

- The regular presence of detectable microbes in pre-eclamptic placentas (e.g. [395; 418; 419])
- The fact that endotoxin (LPS) can act as such a mimic for invoking PE in experimental models
- The fact that every known proteomic biomarker suggested for PE has also been shown to increase during infection.
- The significant number of papers reviewing a link between infection and PE (e.g. [66; 67; 131; 363])
- The almost complete absence (one case) of PE in patients treated with spiramycin [521]

Any and all of these provide powerful strategies for testing whether PE is, in fact, like gastric ulcers [159; 161; 162; 1223], essentially initiated as an infectious disease.

## Footnotes

1 Paper 8 in the series “The dormant blood microbiome in chronic, inflammatory diseases”

## Acknowledgements

BK thanks the Biotechnology and Biological Sciences Research Council (grant BB/L025752/1) for financial support. LCK is a Science Foundation Ireland Principal Investigator (grant number 08/IN.1/B2083). LCK is also The Director of the Science Foundation Ireland-funded INFANT Research Centre (grant no. 12/RC/2272).

